# Intracellular development and impact of a eukaryotic parasite on its zombified microalgal host in the marine plankton

**DOI:** 10.1101/2021.11.04.467241

**Authors:** Johan Decelle, Ehsan Kayal, Estelle Bigeard, Benoit Gallet, Jeremy Bougoure, Peta Clode, Nicole Schieber, Rachel Templin, Elisabeth Hehenberger, Gerard Prensier, Fabien Chevalier, Yannick Schwab, Laure Guillou

## Abstract

Parasites are widespread and diverse in the oceanic plankton, and many of them infect single-celled algae for survival. How these parasites develop and scavenge energy within the host and whether the cellular organization and metabolism of the host is altered remain open questions. Combining quantitative structural and chemical imaging with time-resolved transcriptomics, we unveil dramatic morphological and metabolic changes of the parasite *Amoebophrya* (Syndiniales) during intracellular infection (e.g. 200-fold increase of mitochondrion volume), particularly following digestion of nutrient-rich host chromosomes. Some of these changes are also found in the apicomplexan parasites (e.g. sequential acristate and cristate mitochondrion, switch from glycolysis to TCA), thus underlining key evolutionary-conserved mechanisms. In the algal host, energy-producing organelles (chloroplast) remain intact during most of the infection, but sugar reserves diminish while lipid droplets increase. Thus, rapid infection of the host nucleus could be a zombifying strategy to digest nutrient-rich chromosomes and escape cytoplasmic defense while benefiting from the maintained C-energy production of the host cell.

## Introduction

In aquatic and terrestrial ecosystems, some organisms have developed adaptations to benefit and exploit the metabolism of other organisms through many forms of symbiosis, ranging from commensalism to mutualistic and parasitic interactions. Parasites are important for the functioning and resilience of ecosystems and for the evolution of organisms. While research has largely focused on human and domestic animal parasites, there is a newfound awareness of the relevance of planktonic parasites, particularly in marine ecosystems (Worden et al., 2015). In the past decade, an increasing diversity of eukaryotic parasites has been characterized in the ocean using a combination of DNA sequencing and traditional microscopy, such as Syndiniales, Perkinsozoa and Chytridiomycota (Alacid et al., 2015; Guillou et al., 2008; Jephcott et al., 2016). These parasites are widely distributed in the oligotrophic open ocean and coastal waters (Lima-Mendez et al., 2015; Siano et al., 2011; de Vargas et al., 2015). Several of these parasites infect planktonic microalgae (single-celled photosynthetic eukaryotes), possibly taking advantage of the highly valuable carbon resource produced by the host photosynthetic machinery. Therefore, parasites can regulate algal population dynamics (Coats, 1999), and this is of high ecologically and economically importance when high mortality causes the decline of bloom-forming toxic microalgae in coastal areas (Chambouvet et al., 2008; Montagnes et al., 2008).

The most frequent and diversified marine parasites are Syndiniales (Dinoflagellate), which have a relatively narrow host spectrum. Throughout their evolution, syndiniales have lost their plastids and there is no evidence of a vestigial organelle in their cytoplasm (Farhat et al., 2021), unlike the apicoplast found in apicomplexans (McFadden et al., 1996; Sibbald and Archibald, 2020). While many parasites kill their host before digesting them for their survival, most Syndiniales are biotrophic parasitoids, meaning that their host is maintained alive during most of the infection before cell death (Cai et al., 2020; Jephcott et al., 2016). For instance, the obligate and specialist parasite *Amoebophyra* spp. infects microalgae dinoflagellates (Coats and Park, 2002; Jephcott et al., 2016), which remain photosynthetically active during most of the internal development of the parasite (Kayal et al., 2020). Such infection strategy very likely allows the parasite to efficiently exploit the carbon metabolism of the host (e.g. photosynthetic products) and therefore optimize its growth and replication. While host organelles are physiologically active during infection of *Amoebophyra*, it is not clear how host energy production and carbon storage are impacted by the presence of the parasite. At the end of the infection, the parasite releases numerous motile flagellated zoospores (called dinospores), which do not divide and have a short lifespan (3-15 days) (Coats, 1999; Miller et al., 2012). Syndiniales parasites are therefore strongly dependent on the nutrients and metabolites obtained during their intracellular developmental stages *in hospite* to grow and meet their energy demand while searching for a new host. To date, little mechanistic knowledge is available on the intracellular development of the parasite during infection and its impact on the overall metabolism of its algal host.

Fundamental aspects of the parasitic infection of Syndiniales are still unclear, especially regarding the underlying subcellular mechanisms taking place inside the host cell. To fill this knowledge gap, we used 3-dimensional (3D) electron microscopy combined with transcriptomics to understand the infection strategy of the parasite *Amoebophrya* sp. (strain A120) within its microalgal host (*Scrippsiella acuminata*, Dinophyceae). We particularly investigated the concomitant structural development of the parasite and its impact on the host at the subcellular level. Our approach revealed major morphological and metabolic shifts during the intracellular development of the parasite. By contrast, the bioenergetic machinery of the host is only slightly impacted by the parasite, suggesting that carbon production of the host (starch and lipids) potentially fuels the metabolism of the parasite. Overall, this study provides unprecedented mechanistic insights into a widespread and ecologically important parasite infecting marine phytoplankton. Given that several of these strategies are common to the human parasite apicomplexans, our study also provides new insights into the evolution of parasitism in Alveolata and in eukaryotes more generally.

## Results and Discussion

### Morphological changes of the parasite during its intracellular development

Cultures of the microalga *Scrippsiella acuminata* (dinoflagellate) were infected by the Syndiniales parasite *Amoebophrya* sp. (strain A120). In order to understand the intracellular development within the host, the parasite and its different organelles were reconstructed in 3D after FIB-SEM (Focused Ion Beam Scanning Electron Microscopy) and their volume assessed. Four main morphologically-distinct intracellular stages of the parasite have been observed (Figure 1, Table S1): 1) a transient cytoplasmic parasite (2.2 ± 1.0 μm^3^, n = 4, the only stage surrounded by a parasitophorous vacuole; 2) the young round trophont (44.4 ± 22.1 μm^3^, n = 8) and 3) the mature amoeboid trophont (up to 266 μm^3^), the last two observed in the host nucleus; and 4) the sporont, which occupied most of the host volume. Multiple parasites at the cytoplasmic and trophont stages could be observed simultaneously within a single host cell (Figure S1). During development of these stages, the volume of the parasite increased up to 200-fold with dramatic changes in the morphology and volumes of different organelles such as the mitochondrion, condensed chromatin, and the development of the Golgi apparatus and trichocysts. One of the major morphological changes of the intracellular parasite during infection is the mitochondrion that developed from a small organelle into a reticulate network (Figure 1A-D). Compared to the cytoplasmic parasite (0.040 ± 0.005 μm^3^), the volume of the mitochondrion increased 160 times and 220 times more in the mature trophont (6.4 μm^3^) and sporont (8.8 μm^3^), respectively. Such mitochondrial development in the parasite *Ameobophyra* is similar to the apicomplexan *Plasmodium*, where the mitochondrion forms a branched structure without undergoing fission during some stages of its life cycle (Van Dooren et al., 2005).

**Figure 1:**
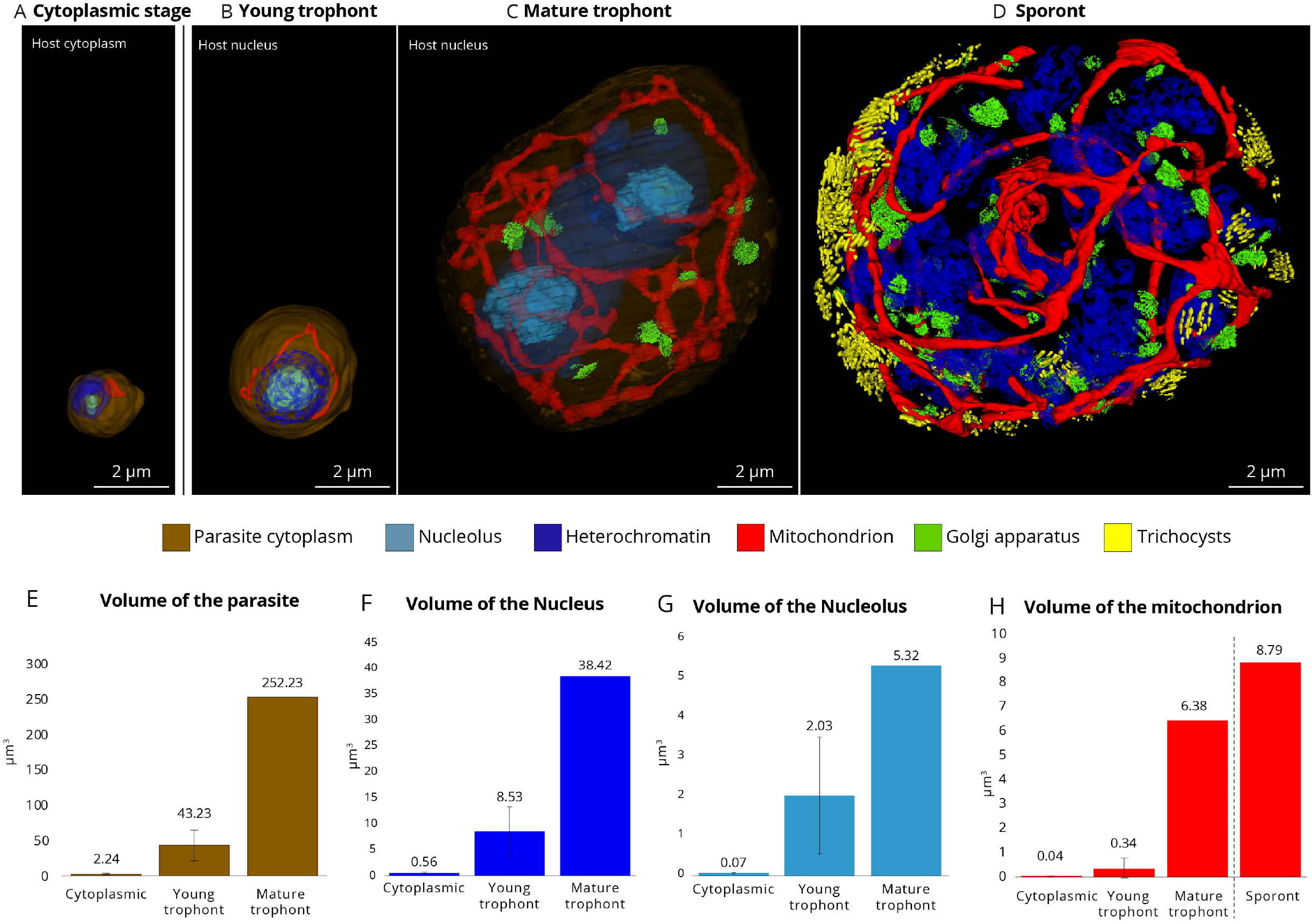
Intracellular development of the marine parasite *Amoebophyra* (Syndiniales) inside its microalgal host (the dinoflagellate *Scrippsiella acuminata*) unveiled by 3D electron microscopy (FIB-SEM: Focused-Ion beam Scanning Electron Microscopy). **A)** 3D reconstruction of the first infection stage (cytoplasmic parasite) in the host cytoplasm where the parasite displayed a relatively small mitochondrion and condensed chromatin at the periphery of the nucleus (Scale bar: 2µm). **B and C)** The parasite then invaded the host nucleus where it developed from a young (B) to a mature trophont (C): the volumes of the parasite, its nucleolus and mitochondrion increased. The Golgi apparatus and dividing nucleus only appeared in the mature trophont. (Scale bar: 2µm). **D)** The sporont parasite exhibited multiple nuclei (without visible nucleolus) and Golgi apparatus, and an extended mitochondrion that occupied the whole parasite volume. Trichocysts were also synthetized at this stage, which are involved in the host attachment for new infection. (Scale bar: 2µm). **E-H)** Volume of the parasite and its organelles (nucleus, nucleolus, mitochondrion) assessed after FIB-SEM-based 3D reconstruction (µm^3^) from four cytoplasmic parasites, seven young trophonts, one mature trophont and one sporont. Brown: parasite volume; light blue: nucleolus; Red: mitochondrion; dark blue: heterochromatin; green: Golgi apparatus; Yellow: trichocysts. See also Table S1 for morphometrics data.

The nucleus and its constituents were also modified during internal development of the parasite. Initially condensed in both dinospores and cytoplasmic parasites, the chromatin was gradually decondensed during the trophont stages, supporting the hypothesis that the early cytoplasmic parasite is a transient stage with minimal transcriptional activity (Figure 1A-B) (Miller et al., 2012). In addition, the volume of the nucleus and nucleolus of the trophont increased by about 13 times and 30 times, respectively, compared to the cytoplasmic stage (Figure 1; Table S1). As ribosome biogenesis is the main function of the nucleolus, we investigated in parallel the expression level of the nuclear ribosomal genes of the parasite in time-resolved transcriptomics data. We found higher expression levels at T24h and T30h (Figure S2), confirming that transcriptional activity and ribosome production significantly increased during the trophont stages. The Golgi apparatus also appeared in the mature trophont and increased in number in the sporont, suggesting that the cellular machinery for protein and lipid production/maturation also occurs at these late stages. The sporont displayed numerous nuclei with peripherally condensed chromatin (lacking visible nucleolus) but without cytokinesis (Figure 1D). This resembles the replication in apicomplexans where daughters cells are formed *de novo* within the cytoplasm without binary fission, along with an elongated mitochondrion (Nishi et al., 2008). Trichocysts that are possibly involved into the attachment of the parasite to a host cell (Miller et al., 2012) were also synthesized at the sporont stage in the parasite *Ameobophyra* (Figure 1D).

After cellular invasion, the host nucleus appeared to be the main cellular target where the parasite settles and accelerates development and metabolism. The 6-fold increase in parasite volume as well as the concomitant growth of the nucleus, mitochondrion and the Golgi apparatus between the two trophont stages demonstrates that significant development of the parasite is accomplished inside the host nucleus. Yet nothing is known regarding the trophic strategy of the parasite nor the fate of the host nucleus and its components (e.g. chromosomes).

### Trophic switch to phagotrophy: Degradation and digestion of host chromosomes

In non-infected cells, the host nucleus contained about 113-119 individual chromosomes (condensed chromatin) with a volume of 0.33 ± 0.10 μm^3^ each (n = 346 chromosomes), representing a total biovolume of 37 ± 2 μm^3^ within the cell (n = 3 cells) (Figure 2A, Table S1). When a young trophont can be detected in the nucleus, the volume of individual host chromosomes can decrease about 3.5 times (in one infected cell: 0.09 μm^3^ ± 0.04 μm^3^, n = 125 chromosomes) (Figures 2B-2F, Table S1). We also observed a decrease in the volume of the host nucleolus from 5.1 ± 0.9 μm^3^ in non-infected hosts (n = 3 cells) down to 2.7-1.9 μm^3^ in infected hosts, suggesting diminished ribosome production that may lead to lower transcriptional activity (Figure 2G, Table S1). This can explain the steady decline in the number of total host transcripts described previously (Kayal et al., 2020). The degradation of the genetic material of the host can potentially be triggered by the parasite. However, we were unable to unambiguously identify genetic signatures related to extracellular chromosome degradation (e.g. nucleases participating in the purine and pyrimidine metabolism) in the genome of the *Amoebophrya* parasite. Nevertheless, given that the young trophont parasite is surrounded by an intact and relatively thick membrane with no cytoplasmic invagination, we suggest that importation of nutrient and metabolites in the first stages of its intranuclear development could only be through osmotrophy.

**Figure 2:**
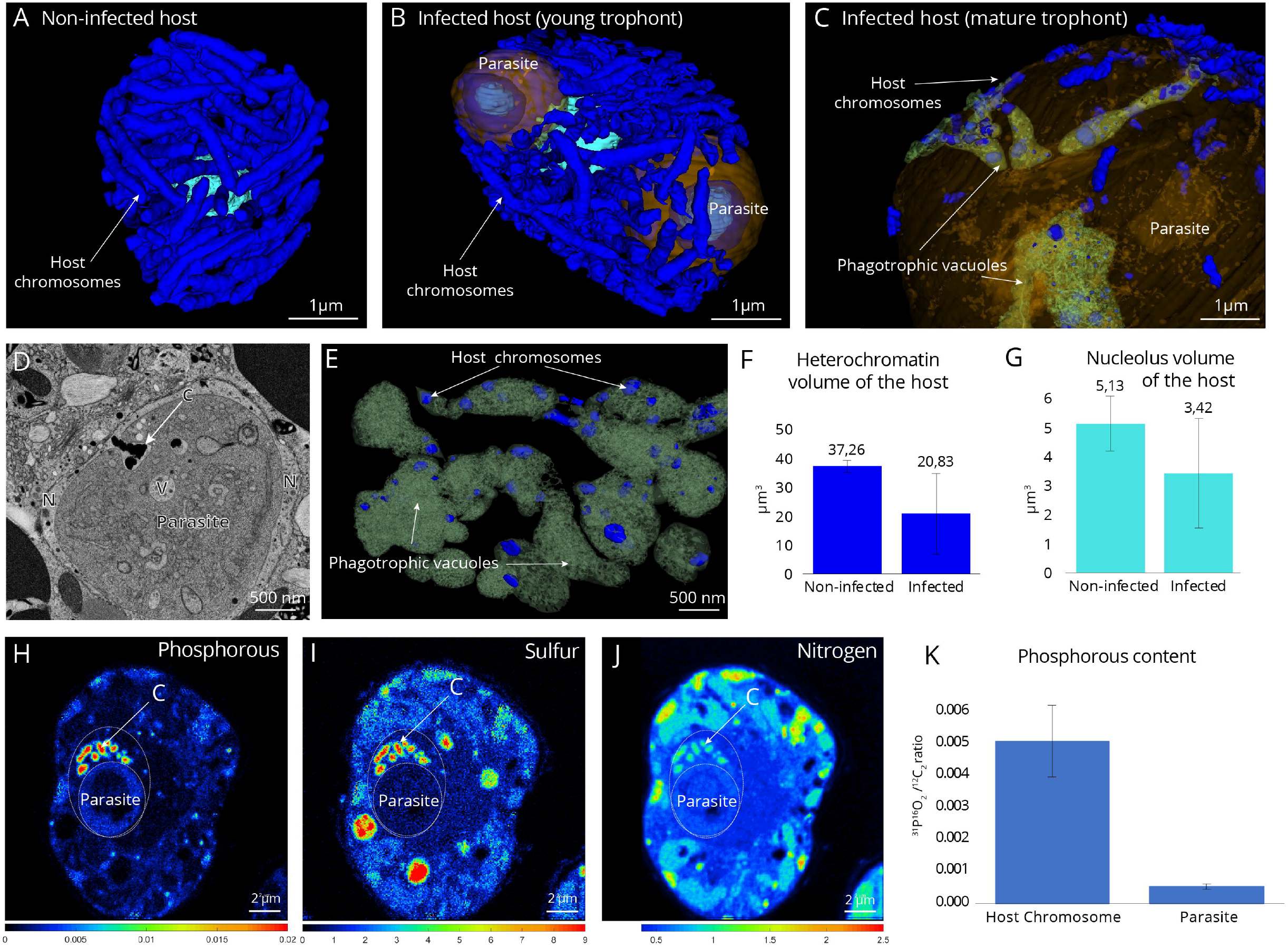
Degradation and digestion of the host chromosomes and nucleus by the parasite *Amoebophyra* unveiled by 3D electron microscopy and nanoSIMS. **A)** 3D reconstruction of the nucleolus and individual chromosomes of non-infected hosts (about 113-119 per host cell of about 0.33 ± 0.10μm^3^ each; n = 346). **B)** Host nucleus, infected by two trophont parasites, displayed smaller chromosomes and nucleolus compared to non-infected hosts. **C - E)** At later infection stages, the mature trophont parasite developed multiple phagotrophic vacuoles to engulf and ingest host chromosomes. D) Transmission electron microscopy (TEM) micrograph showing the engulfment of an electron-dense host chromosome (C) into the vacuole (V) of a mature trophont parasite within the host nucleus (N). **F and G)** Volumes of the heterochromatin and nucleolus (in μm^3^) of non-infected and infected host cells assessed after FIB-SEM-based 3D reconstruction. **H-J)** NanoSIMS (Nanoscale Secondary Ion Mass Spectrometry) mapping of the elements Phosphorous (H, ^31^P^16^O_2_/^12^C_2_), Sulfur (I, ^34^S/^12^C_2_) and Nitrogen (J, ^12^C^14^N/^12^C_2_), showing that host chromosomes (C) are highly concentrated in these nutrients compared to the nuclear parasite. (Scale bar: 2µm). **K)** Phosphorous (P) content calculated as ^31^P^16^O_2_/^12^C_2_ from nanoSIMS ion count map in the host chromosomes and parasite cell (including nucleus and cytoplasm). P content of the host chromosomes (n = 131) were estimated to be about 10 times more important than in the parasite cell (n = 22) (See also Table S3). Brown: parasite volume; light blue: nucleolus; Red: mitochondrion; dark blue: heterochromatin; green: Golgi apparatus; Yellow: trichocysts. See also Table S1 for morphometrics data.

By contrast, in the mature trophont, multiple phagotrophic vesicles were observed indicating a switch of the trophic mode from osmotrophy to phagotrophy (Figure 2C-E). The 3D reconstructions revealed invaginations of the cytoplasmic membrane at several locations creating a tubular network of vacuoles (previously described as a cytopharynx (Cachon, 1964)), in which degraded chromosomes of the host (0.03 ± 0.01 μm^3^) were engulfed, vascularized and digested. In order to further understand this trophic switch, we investigated the expression levels of genes encoding key proteins involved in formation and acidification of vacuoles and phagocytosis (Cox et al., 2000; Harris et al., 2001; Marshansky et al., 2014; Vines and King, 2019): subunits of the vacuolar H^+^-ATPase (V-ATPase) and the GTPase Ras-related protein Rab-11. We found that Rab11 and V-ATPase subunits exhibited no expression in dinospores but maximum expression at T24h and T30h-T36h, respectively, corresponding to late intracellular stages (i.e. trophont and sporont) (Figure S3, Table S2). We therefore hypothesize that Rab11 expression could be related to the formation of phagotrophic vacuoles where V-ATPase activity would generate an acidic environment for chromosome digestion.

Subcellular nutrient mapping by NanoSIMS (Nanoscale Secondary Ion Mass Spectrometry) imaging on cellular sections showed that the chromosomes of the host were highly concentrated in phosphorous (P), sulfur (S), and nitrogen (N), representing a concentration hotspot of these nutrients in the host cell (Figures 2H-K, and S4). For instance, P (^31^P^16^O_2_/^12^C_2_) and S (^34^S/^12^C_2_) content of the host chromosomes (n = 131) were estimated to be about 9.8 and 8.5 times more than values measured in the parasite cell (n = 22), respectively (Table S3). Similarly, N content (^12^C^14^N/^12^C_2_) in the host chromosomes is about 1.8 times more than in the parasite cell. Thus, although nutrient transfer cannot be unambiguously demonstrated here, we hypothesize that the parasite benefits from the degradation and digestion of nutrient-rich host chromosomes. DNA can also be a valuable source of carbon for the parasite (Pinchuk et al., 2008). Rapid infection of the host nucleus therefore appears to be a key strategy to have direct access to major nutritional resources from the host required for growth and replication (e.g. C, N, P) while escaping cytoplasmic host defense. In line with this hypothesis, the substantial increase of the volumes of the parasite and its developing organelles (e.g. nucleus, mitochondrion, and the Golgi apparatus) (Figure 1C) clearly reflects a rewiring of parasite metabolism and growth strategy during the phagotrophic stage.

During this phagotrophic stage, we also observed a significant development of a network of tubules in the host nucleus, which highly resembles the Intravacuolar Network (IVN) described in the human parasite *Toxoplasma* (Caffaro and Boothroyd, 2011; Lopez et al., 2015). During *Ameobophyra* infection, the IVN-like structures often surrounded and concentrated around the host chromosomes (Figure S5). IVN has been proposed to be involved in nutrient and lipid uptake in *Toxoplasma gondii* (Nolan et al., 2017; Pszenny et al., 2016) and could play the same role here in this planktonic parasite. Yet, we were not able to identify homologs in the *Amoebophrya* genome of the two key dense granule proteins genes, GRA2 and GRA6, responsible for shaping the IVN of *T. gondii* (Fox et al., 2019; Lopez et al., 2015), possibly because of highly divergent sequences not identifiable by homology (Farhat et al., 2021).

In addition to the digestion of the chromosomes as a putative nutritional resource, intracellular parasites may also rely on the host central carbon metabolism for powering their development and replication to successfully produce infectious free-living dinospores. We therefore investigated whether the carbon metabolism and storage capacity of the host were remodeled during the infection and identify potential carbon sources for the parasite.

### Impact of the parasitic infection on the host bioenergetics

3D electron microscopy allowed us to reconstruct and quantify organelles and compartments of the algal host that are central for its bioenergetics before (non-infected hosts) and during the parasitic infection. We particularly focused on the cellular sites for carbon fixation (plastids and pyrenoid) and storage (lipids and starch) in order to assess the impact of the parasite on the central carbon metabolism and carbon partitioning of its host. While the volume occupancy of the mitochondria was similar between non-infected and infected host cells (4.9% and 4.5% respectively, Table S1), the plastid occupancy only slightly decreased during infection from 17.2 ± 3.2 % of the cell volume in non-infected (n = 3 cells) to 14.0 ± 1 % in infected hosts (n = 3 cells) (Figure 3, Table S1). In addition, the overall structure of the plastids remained intact with similar arrangement of thylakoid membranes (Figure S1). Although the number of pyrenoids - rubisco-containing compartment where CO_2_ is fixed (Freeman Rosenzweig et al., 2017) - varied in host cells, the pyrenoid volume occupancy remained stable in the host cell throughout the infection (1.7% of the cell volume in both infected and non-infected host cells). The maintenance of the volumes and morphology of plastid and pyrenoid indicates that the host capability for carbon fixation is not impacted by the parasites during the first stages of infection, and sugar production is potentially maintained. These results corroborate a previous study that showed stable quantum yield of photosystem II (F_v_/F_m_) and plastid pigments content, along with continuous expression of chloroplast-encoded photosynthetic genes during most of the infection (Kayal et al., 2020). It is interesting to note that dinospore production was 5-fold lower in darkness compared to light conditions (Kayal et al., 2020), stressing the importance of the host photosynthetic machinery and energy production for successful parasite replication.

**Figure 3:**
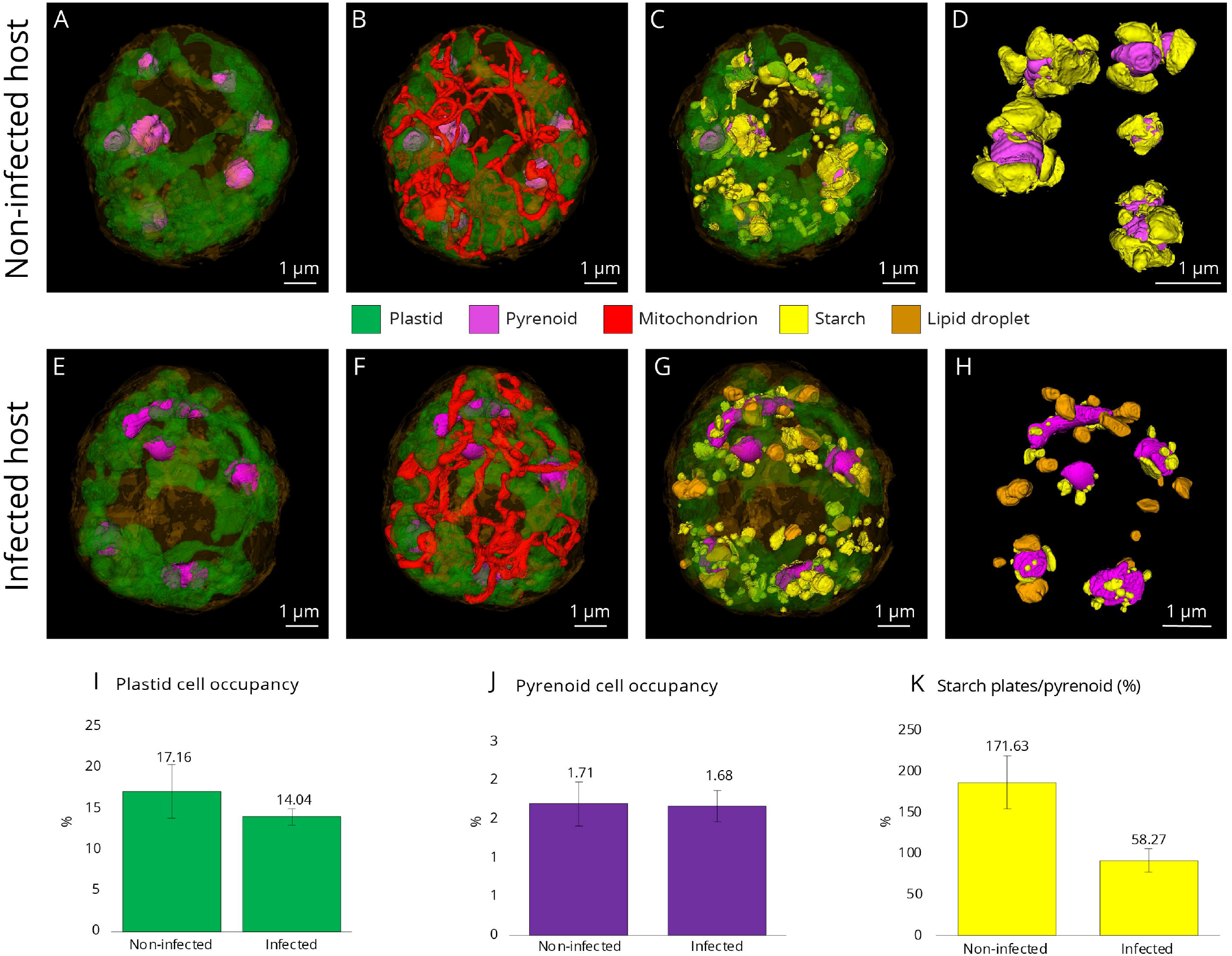
3D cellular architecture of non-infected and infected microalgal host cells (the dinoflagellate *Scrippsiella acuminata*) unveiled by FIB-SEM with a focus on the bioenergetic machinery and carbon reserves. **A-D)** 3D reconstruction of the non-infected host cells with A) its plastid (green) and C-fixing pyrenoids (purple), B) mitochondrion; C) starch grains and plates (yellow); and D) starch plates (yellow) around the pyrenoids (purple). **E-H)** 3D reconstruction of the infected host cells with E) its plastid (green) and pyrenoids (purple), F) mitochondrion; G-H) Starch (yellow) and lipids (orange). **I-K)** Volume occupancy (% of the cell volume) of the plastid (I) and the pyrenoid (J), and the volume ratio between the starch plates and the pyrenoid (K) in three non-infected and three infected hosts cells after FIB-SEM-based 3D reconstruction.

To better evaluate carbon storage in non-infected and infected hosts (cryofixed at the same time), we quantified the volume of starch, which is a semicrystalline form of polysaccharides storage located in the cytoplasm in the form of grains and attached to the pyrenoid as plates (Figures 3C-3G). The total starch occupancy (grains and plates; reconstructed in 3D) decreased in infected cells, changing from 5% of the cell volume before infection to 3% during infection (Table S1). Starch was even absent in one host cell infected by three trophont parasites and almost absent at the final sporulation stage. This overall decrease in starch volume is mainly explained by the two-fold decrease in the volume of the starch plates around the pyrenoid. More particularly, starch plates represented on average 171.6% of the pyrenoid volume in non-infected cells (i.e. starch plates were 1.75 times more voluminous than the pyrenoid) and 58.2% during the infection (i.e. two times less voluminous than pyrenoid) (Figures 3D; 3H and 3K). Such decrease in host sugar reserve suggests two different but potentially concomitant scenarios: the host machinery for sugar production is 1) functional but is slowed down during the infection, and/or is 2) functional but starch is rapidly consumed by host and parasite and/or rewired into the central carbon metabolism of the host. The presence of several host transcripts for cytosolic soluble and granule-bound starch synthases at each infection stage indicates the potential for starch synthesis throughout the infection (Table S4). The host also encodes for plant homologs involved in initial starch mobilization (a cytosolic alpha-glucan and phosphoglucan water dikinase) and enzymes degrading the mobilized starch (beta-amylases and isoamylases) (Zeeman et al., 2010), thus potentially providing soluble sugars to the parasite. Transcripts for these host enzymes were found present until the last stage of infection (Table S4). In contrast, we were unable to identify any of those starch degradation genes in the genome of the parasite. It is therefore possible that the parasite scavenges sugar molecules, such as glucose, directly from the host.

The growth and replication of apicomplexan parasites rely on a continuous supply of host-derived sugar via different transporters (Qureshi et al., 2020). For example, the hexose transporter PfHT1 can transport both glucose and fructose across the cell membrane of *Plasmodium falciparum* (Blume et al., 2009; Qureshi et al., 2020). We therefore searched for two families of sugar transporters in the genome of *Amoebophrya* (strain A120): hexose transporters (HT) and Sugars Will Eventually be Exported Transporters (SWEET). Using similarity searches and phylogenetic analyses, we identified one SWEET-like protein predicted to have at least six predicted transmembrane domains (TMs), as well as three hexose transporters (HTs) displaying 11-12 predicted TMs (Figures S6 and S7). We then investigated their expression level at different stages of the infection and compared them to the dinospore stage. The SWEET gene and one HT (HT1) gene had maximum expression during the intracellular stage (T30h-T36h), with nearly no expression in dinospores (Figure 4A). We hypothesize that SWEET and HT1 are likely involved in sugar scavenging from the host during the intracellular development of the parasite. SWEETs are known for bidirectional passive transport of various mono- and disaccharides from high to low sugar concentrations (Chen et al., 2012; Latorraca et al., 2017). High concentration of sugars in the host could allow the parasite to “passively” obtain these metabolites through its SWEET transporter without energy consumption. The two other HT genes (HT2 and HT3) were mainly expressed at later intracellular stages (T36h) and in dinospores, suggesting a sequential role of these transporters for sugar transport within the cell during the life cycle of the parasite (Figure 4A).

**Figure 4:**
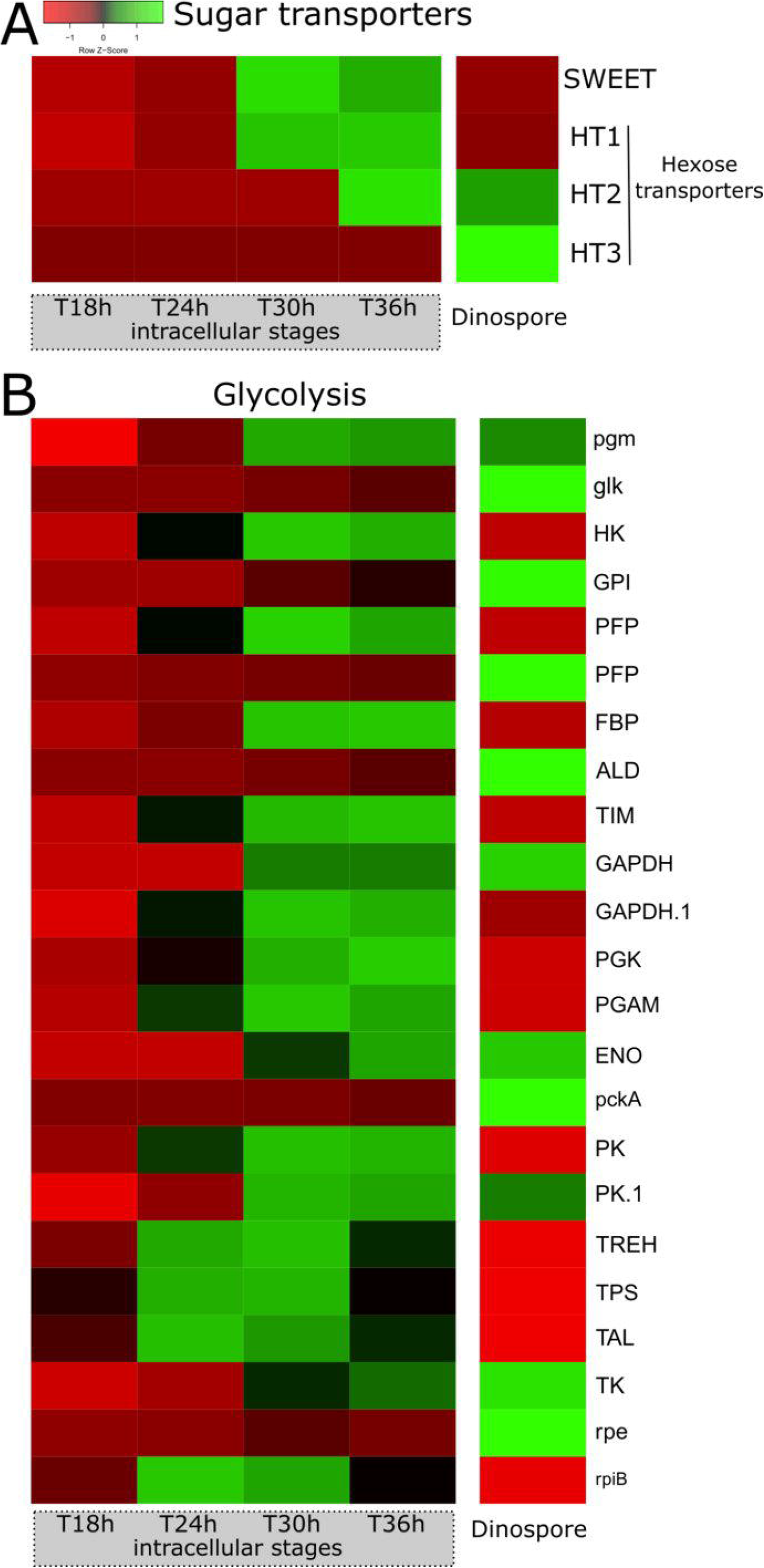
Expression levels of genes involved in sugar transport and glycolysis of the marine parasite *Amoebophyra* across different intracellular stages within its host (the dinoflagellate *Scrippsiella acuminata*) and in dinospores (extracellular). **A)** Heatmap showing the expression level of four genes of the parasite encoding putative sugar transporters during the infection (T18h, T24h, T30h and T36h) and the dinospore stage: one SWEET (Sugars Will Eventually be Exported Transporters) and three hexose transporters (HT1, HT2, and HT3). (See also Figures S6, S7 and S9, Table S2). **B)** Heatmap showing the expression level of genes of the glycolysis pathway of the parasite during the infection (T18h, T24h, T30h and T36h) and in dinospores. The list of genes, their sequences and expression values can be found in Table S2.

Lipid droplets are also a major carbon storage in microalgae. These reserves contain neutral lipids such as cholesterol esters and triacylglycerols (TAG) and have been shown to be key players in host-pathogen interactions (Nolan et al., 2017). In electron microscopy, lipid droplets are recognizable as homogeneous electron-dense structures without a membranous bilayer. Contrary to non-infected host cells, we observed large lipid droplets in the cytoplasm of infected hosts, representing a volume from 19.4 μm^3^ up to 49 μm^3^ (between 1.33% and 2.3% of the host volume) (Figures 3G and 3H). While most lipid droplets were closely associated with the plastid and mitochondrion, some were attached to the host nucleus (Figure S8). Host transcripts for the complete FASII fatty acid biosynthesis pathway, which provides fatty acids for incorporation into TAGs, were detected throughout the infection, as well as several isoforms of diacylglycerol acyltransferase (DGAT; catalyzing the last step of TAG formation), and acetyl-CoA acetyltransferase (ACAT; involved in cholesterol esterification) (Table S4). Increase in host lipid droplets was also observed during the infection of *Toxoplasma* and *Plasmodium* (Amiar et al., 2020; Hu et al., 2017; Nolan et al., 2017) and represents a lipid scavenging strategy (Gomes et al., 2014; Hu et al., 2017; Nolan et al., 2017). Future investigations to see whether and how the *Amoebophyra* parasite benefits from this production of lipids in the host will shed light on its metabolic strategy during infection and evolutionary-conserved strategies in parasitic alveolates across ecosystems and hosts.

Here, we provide evidence that the photosynthetic machinery and the carbon metabolism of the zombified host are still active but altered during infection by *Ameobophyra*. The associated production of sugars and lipids could represent an energetic source to fuel the metabolism of the parasite (notably for ATP production). An evaluation of the metabolic activity of the parasite is therefore of primary importance to fully understanding its life cycle and adaptations to thrive within the intracellular host environment.

### Metabolism of the intracellular parasite (ATP production)

Cellular ATP can be produced by cytoplasmic glycolysis, as well as the mitochondrial Tricarboxylic acid (TCA) cycle and oxidative phosphorylation (OXPHOS) pathway. We reconstructed the glycolysis pathway in *Amoebophyra* and assessed the expression levels of its constituent genes at the intracellular and dinospore stages using time-resolved transcriptomics data (Figures 4B and S9). Overall, we observed distinct expression patterns where genes of the preparatory phase of glycolysis (consumption of ATP) were expressed in both intracellular and dinospore stage while most genes of the pay-off phase (production of ATP and NAD(P)H) were expressed during intracellular infection (Figures 4B and 6). Overall, gene expression analysis suggests an active glycolysis of the intracellular parasite, which leads to the production of pyruvate that can potentially fuel the TCA cycle. However, we found that all the genes involved in the TCA cycle were mostly expressed in dinospores (Figures 5A and 6). Similarly, the Mitochondrial Pyruvate Carrier that allows the pyruvate to enter the mitochondria was also only expressed in dinospores (Figure 6). Therefore, these results imply that there is no complete oxidation of carbohydrates and lipids during infection within the host, and the intracellular parasite does not rely on the TCA cycle to produce ATP (and NAD(P)H). This is also the case during the asexual stage of *Plasmodium* that mainly relies on glycolysis to produce ATP, while TCA metabolism occurs at low turnover (Jacot et al., 2016). This metabolic strategy is found in highly proliferating cells (e.g. apicomplexans, cancer cells) that undergo low respiration and increased glycolysis to support biomass generation with glycolytic intermediates (also known as the Warburg effect) (Salcedo-Sora et al., 2014). Such high glycolytic flux typically occurs in glucose-replete environments, which is likely the case here for the parasite *Ameobophyra* inside its physiologically active algal host.

**Figure 5:**
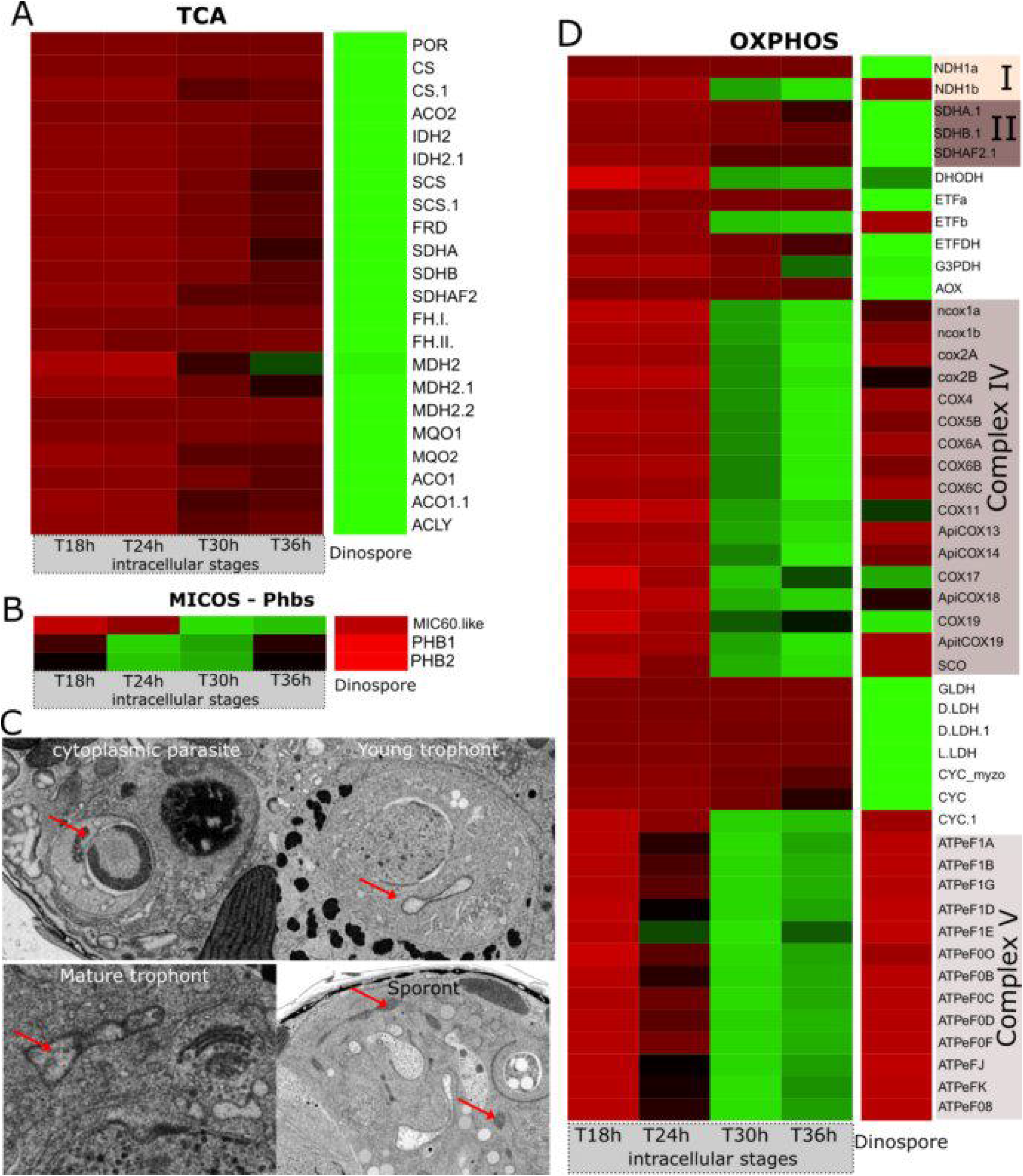
Expression levels of genes involved in mitochondrial respiration and formation of cristae in the parasite *Amoebophyra* across different intracellular stages within its host (the dinoflagellate *Scrippsiella acuminata*) and in dinospores (extracellular). **A and D)** Heatmap showing the expression level of genes of the TCA cycle (A) and the OXPHOS (D) pathway of the parasite. See also Figure S10 and Table S2; **B)** Expression levels of genes encoding MiC60 from the MICOS complex (MItochondrial contact site and Cristae Organizing System), and Phb1 and Phb2 genes encoding prohibitin ring complexes. **C)** Transmission electron microscopy (TEM) micrographs showing the internal morphology of the mitochondrion of the parasite at different infection stages. In the cytoplasmic parasite, the electron dense mitochondrion harbored cristae (internal invagination of the inner mitochondrial membrane), which were absent in the mitochondrion of the nuclear trophont parasites (young and mature trophonts). Some vesicles could be observed in the mitochondrion of the mature trophont. Cristae reappeared in the sporont stage where the mitochondrion was substantially developed. The list of genes, their sequences and expression values can be found in Table S2.

**Figure 6:**
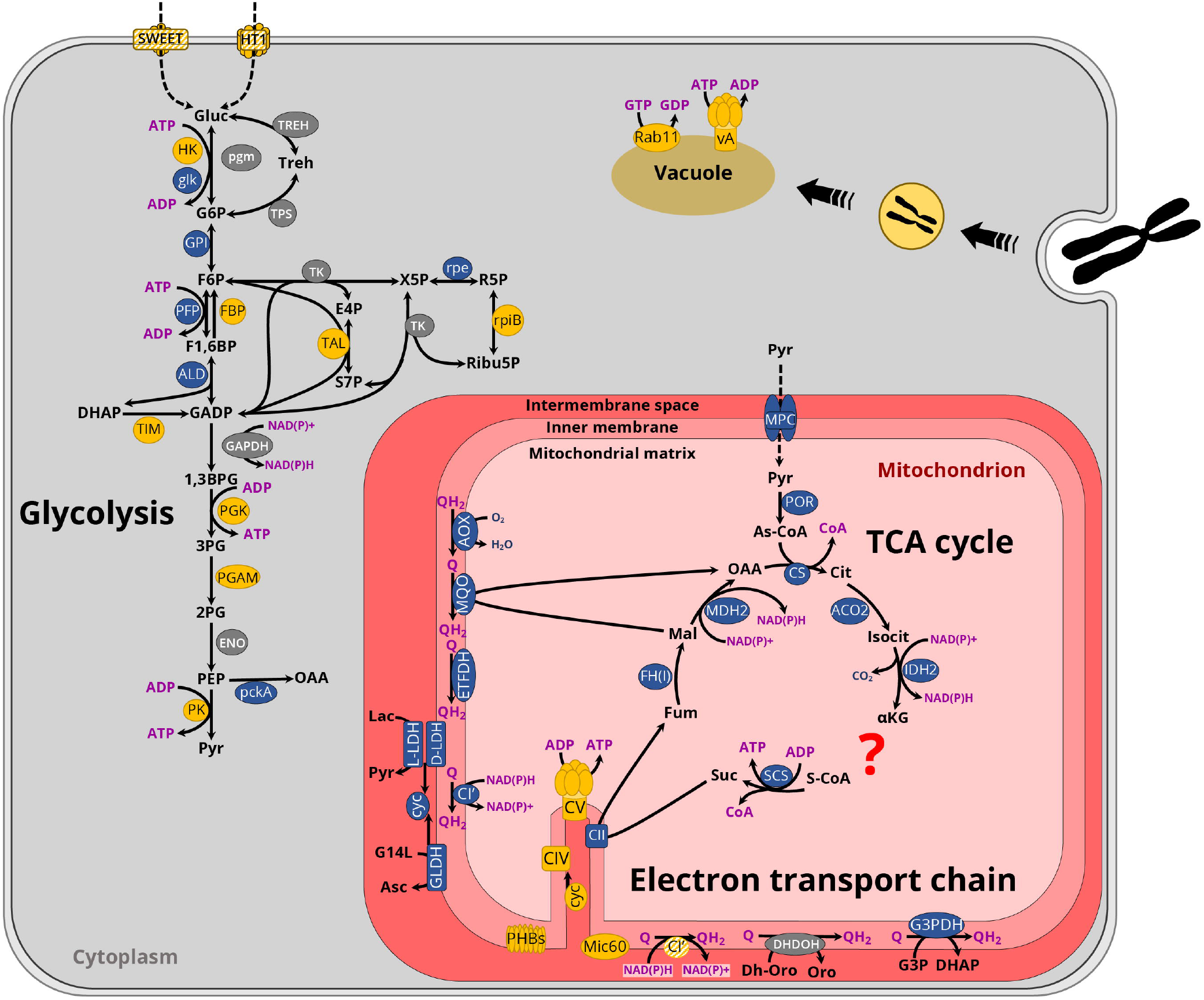
Schematic overview of the intracellular metabolism of the marine parasite *Amoebophyra* inside its microalgal phototrophic dinoflagellate host *Scrippsiella acuminata* underlying major metabolic shifts. Major energy pathways have been displayed where the color of individual enzymes reflects the maximum expression of their genes: orange for intracellular parasites; blue in dinospores; grey suggesting no difference in expression between the two stages. Dashed lines represent transport of various components; filled lines represent enzymatic reactions. Coenzymes are color-coded as purple: adenosine bi- and triphosphate (ADP and ATP, respectively); nicotinamide adenine dinucleotide phosphate (NAD(P)H); coenzyme A (CoA); quinone pool (Q and QH_2_). The putative location of one of the NADH:ubiquinone oxidoreductase (NDH1b/CI’) complexes, the hexose transporter (HT1) and Sugars Will Eventually be Exported Transporter (SWEET) are depicted by colored hatchings. A question mark in the tricarboxylic acid cycle (TCA) illustrates the apparent loss of the oxoglutarate dehydrogenase/ (OGDC) or α-ketoglutarate dehydrogenase complex in *Amoebophrya*. Invagination of the cytoplasm to form a cytopharynx is represented to illustrate captured host chromosome to be digested in vacuoles. The list of genes, their sequences and expression values can be found in Table S2.

Reliance on glycolysis and low respiration led to the assumption that the mitochondrion of the parasite *Amoebophyra* might be metabolically quiescent in the intracellular stage. Yet, we observed a substantial development of the mitochondrion during the infection (220-fold increase of the volume) (Figure 1). To further understand mitochondrial activity, we investigated the internal morphology as well as the expression level of genes involved in the OXPHOS pathway. The single electron-dense mitochondrion in cytoplasmic parasites displayed typical cristae (internal invagination of the inner mitochondrial membrane) that are also found in the free-living dinospore (Miller et al., 2012) (Figure 5C). Then, the mitochondrion developed as an empty “tube” without forming cristae in the nuclear young trophont. In the mature trophont, vesicle-like structures were observed in the mitochondrion (Figure 5C). Reappearance of canonical cristae only occurred during the sporont stage within the reticulate mitochondrion. This is similar to the development of the mitochondria of *Plasmodium* whereby cristae are also temporarily absent in asexual blood-stage and reform in the sexual stage (gametocytes) (Evers et al., 2021; Krungkrai, 2004; Prapunwattana and Krungkrai, 1999).

Cristae play a central role in cellular metabolism since they are sites of high concentration of protons (protonic capacitance) where ATP is generated (Lee, 2020). In eukaryotes, cristae formation and stabilization mainly rely on the interplay of three players, namely the MItochondrial contact site and Cristae Organizing System (MICOS), the large GTPase optic atrophy 1 (OPA1), and the mitochondrial ATP synthase (F_1_F_0_- ATPase, also known as complex V of OXPHOS) (Pánek et al., 2020; Stephan et al., 2020). The presence of cristae is also concomitant with prohibitin ring complexes encoded by Phb1 and Phb2 genes (Wideman and Muñoz-gómez, 2016). In *Toxoplasma*, the assembly of ATP synthases into hexamers was also shown to be responsible for cristae invagination (Mühleip et al., 2021). By investigating the genome of the parasite *Ameobophyra*, we identified two prohibitin genes (Phb1 and Phb2) but only found one gene (Mic60) of the MICOS complex and no homolog for OPA1. Transcriptomics data revealed similar expression pattern for Mic60 and components of complex V, both displaying maximum expression at T30h-T36h and low or no expression in the early infection stages and in dinospores (Figures 5B and 5D). Similarly, the prohibitin genes Phb1 and Phb2 were mainly expressed during the intracellular phase with nearly no expression in dinospores (Figure 5B). Thus, gene expression levels suggest that cristae formation and stabilization occur in late intracellular stages and explain acristate mitochondrion observed in young trophonts (Figure 5C).

Cristae formation can be linked to the production of ATP through the dissipation of the proton gradient generated by the mitochondrial Electron Transport Chain (ETC). The OXPHOS pathway in *Amoebophrya* is divided into two independently operating subchains (Farhat et al., 2021), very similar to what has been described in *Chromera velia* (Flegontov et al., 2015) (Figure S10). In the parasite *Ameobophyra*, the proton gradient is only generated by the cytochrome *c* oxidase (complex IV) and is dissipated by complex V (Farhat et al., 2021; John et al., 2019). We found that genes of complexes IV and complex V displayed maximum expression at T30h and T36h, along with one homolog of cytochrome C (CYC) and NADH:ubiquinone oxidoreductase (NDH1 or CI’) (Figures 5D and 6, and S10). Such expression pattern suggests that ETC-based ATP production starts to occur during the late intracellular stages of the infection which coincides with the formation of cristae at the sporont stage. These results also show that many components of the mitochondrial OXPHOS are dispensable during the intracellular stage, except the Dihydroorotate dehydrogenase (DHODH) for pyrimidine synthesis (Figure S10), as found in the malaria parasites (Painter et al., 2007).

By contrast, in the dinospore stage, the concomitant expression of all genes of the TCA cycle (including the Mitochondrial Pyruvate Carrier) and most of OXPHOS pathway (excluding complexes IV and V) reveals that the parasite maintains an active catabolism while searching for a new host (Figures 5 and 6, and S10). Although it is not known whether the parasite can feed *ex hospite*, we hypothesize that the lifespan of the free-living dinospores and the success rate of new infections strongly depend on carbon reserves accumulated during the intracellular stage from the host bioenergetics (e.g. sugar, lipids) that can fuel TCA and OXPHOS in the glucose-deplete oceanic waters.

## Conclusion

Combining transcriptomics and 3D electron microscopy, we shed light on the intracellular development of the marine parasite *Ameobophyra* within its microalgal host and unveiled dramatic shifts in trophic strategy and metabolic activity. Upon host entry, the cytoplasmic parasite appears to be metabolically and transcriptionally quiescent until it invades the host nucleus. Then, the growing trophont undergoes major morphological changes, particularly following the trophic switch into phagotrophy where host chromosomes are engulfed and digested. We presume that this trophic strategy provides a highly valuable source of carbon and nutrients for the development and growth of the parasite. The intracellular parasite might also benefit from the host sugar production to fuel glycolysis for ATP production. This is possible since the zombified host is photosynthetically active with preserved plastids and pyrenoids such that production of sugar is maintained during infection. It is also possible that a lipid scavenging strategy takes place to benefit from the lipid droplets of the host that increase during infection. ATP production in the intracellular parasite can also occur at late stages through the activity of mitochondrial complexes IV and V. This can be reflected by the morphological plasticity of the mitochondrion which first expands without cristae in the nuclear parasites. Mitochondrial cristae reappear at the sporont stage before the expanded mitochondrion is distributed to each dinospore that requires higher respiration. Indeed, we provide evidence that the free-living parasite switches the energy metabolism towards oxidative phosphorylation and may rely on carbon reserves salvaged and accumulated during the intracellular development.

It is interesting to note that the intracellular development and metabolism of the planktonic parasite *Ameobophyra* living in marine ecosystems exhibit many similarities with the apicomplexans *Toxoplasma and Plasmodium*. These alveolate parasites also have a highly flexible mitochondrial development and display sequential acristate and cristate mitochondria during their intracellular life cycle (Krungkrai, 2004; Voleman and Doležal, 2019). This can be linked to their metabolic strategy with high glycolytic activity and low respiration at one particular stage of infection. In addition, we also observed the IVN-like network, synthesis of lipid droplets in the host upon infection and phagotrophic activity to ingest host material (Elliott et al., 2008; Fox et al., 2019; Nolan et al., 2017). These mechanisms appear to be evolutionarily conserved across parasitic alveolates in different ecosystems regardless of their eukaryotic hosts, thus underlining their importance during infection. Future investigations are required to study the role of the IVN-like network as well as the molecular players that are involved in the sugar and lipid scavenging in this marine parasite. Metabolomics will also have the potential to improve our understanding of the metabolic rewiring of the parasite during infection and identify potential metabolites scavenged from the host. This knowledge will be essential to elucidate the survival of the free-living dinospores and understand the ecological success of this widespread parasite that has a major impact on phytoplankton populations and therefore carbon cycling in the ocean.

## Methods

### Culture conditions

The dinoflagellate *Scrippsiella acuminata* ST147 (RCC 1627) was maintained in F2 medium (enriched with 5% of soil v/v) in these following culture conditions: 20°C, 80-100 μmol photons m^-2^s^-1^, L:D cycle of 12:12 h. (More information on the culture provided here: dx.doi.org/10.17504/protocols.io.vrye57w). The parasite *Amoebophrya ceratii* A120 (RCC 4398) (syndiniales, amoebophryidae, equivalent to Marine ALVeolates Group II, or MALV-II) was maintained by inoculating frequently (every 3-4 days) fresh cultures of *Scrippsiella acuminata* (3-4 days old). For the infection experiment, dinospores (*ex hospite*) of the parasite *Amoebophrya* were obtained by filtering by gravity on a 5µm mesh size polycarbonate filter and were mixed with a fresh culture of host cells (1 vol dinospore cells for 2 volumes of host cells) for 35 hours.

### 2D and 3D Electron microscopy (TEM and FIB SEM)

#### Sample preparation

The non-infected and infected microalgae *Scrippsiella* were concentrated on a 5 μm mesh size polycarbonate filter. Cells were then collected and cryo-fixed using high-pressure freezing (HPM100, Leica), followed by freeze-substitution (EM ASF2, Leica) as in (Decelle et al., 2019; Uwizeye et al., 2021a). For the freeze substitution (FS), a mixture 2% (w/v) osmium tetroxide and 0.5% (w/v) uranyl acetate in dried acetone was used for FIB-SEM with a programed protocol from (Decelle et al., 2019). For TEM and nanoSIMS, the FS mix contained only 1% of osmium tetroxide For TEM analysis, ultrathin sections of 60 nm thickness were mounted onto copper grids or slots coated with formvar and carbon. Sections were then stained in 1% uranyl acetate (10 min) and lead citrate (5 min). Micrographs were obtained using a Tecnai G2 Spirit BioTwin microscope (FEI) operating at 120 kV with an Orius SC1000 CCD camera (Gatan).

#### FIB-SEM acquisition

Samples were mounted onto the edge of a SEM stub (Agar Scientific) using silver conductive epoxy (CircuitWorks) with the trimmed surfaces facing up and towards the edge of the stub. Samples were gold sputter coated (Quorum Q150RS; 180 s at 30 mA) and placed into the FIB-SEM for acquisition (Crossbeam 540, Carl Zeiss Microscopy GmbH). Atlas3D software (Fibics Inc. and Carl Zeiss Microscopy GmbH) was used to perform sample preparation and 3D acquisitions. First, a 1 μm platinum protective coat (20-30 μm^2^ depending on ROI) was deposited with a 1.5 nA FIB current. The rough trench was then milled to expose the imaging cross-section with a 15 nA FIB current, followed by a polish at 7 nA. The 3D acquisition milling was done with a 1.5 nA FIB current. For SEM imaging, the beam was operated at 1.5 kV/700 pA in analytic mode using an EsB detector (1.1 kV collector voltage) at a dwell time of 8 μs with no line averaging. For each slice, a thickness of 8□or 10 nm was removed, and the SEM images were recorded with a pixel size of 8 or 10□nm, providing an isotropic voxel size of 512 nm^3^ or 1000 nm^3^. Raw electron microscopy data are deposited in EMPIAR, accession code EMPIAR-XXXXX.

#### 3D reconstruction and volume quantification

From the stack of images, regions of interest were cropped using the open software Fiji (https://imagej.net/Fiji), followed by image registration (stack alignment), noise reduction, semi-automatic segmentation, 3D reconstruction of cells and morphometric analysis as described previously (Uwizeye et al., 2021b). Image registration was done by the FIJI plugin ‘Linear Stack Alignment with SIFT’(Lowe, 2004), then fine-tuned by AMST (Hennies et al., 2020). Aligned image stacks were filtered to remove noise and highlight contours using a Mean filter in Fiji (0.5-pixel radius). Segmentation of organelles (plastids, mitochondrion, nucleus) and other cellular compartments of the parasite and the host cells (starch, lipid) was carried out with 3D Slicer software (Kikinis et al., 2014) (www.slicer.org), using a manually-curated, semi-automatic pixel clustering mode (5 to 10 slices are segmented simultaneously in z). We assigned colors to segmented regions using paint tools and adjusted the threshold range for image intensity values. Morphometric analyses were performed with the 3D slicer module “segmentStatistics” on the different segments (segmented organelles) and converted to μm^3^ considering the voxel size of 512 or 1000 nm^3^ (Table S1). In total, we analyzed three non-infected host cells, six infected host cells, and 13 parasites.

### NanoSIMS measurements

Semi-thin sections (200-300 nm) on silicon wafers were coated with 20-nm gold-palladium and analyzed with a nanoSIMS 50L (Cameca, Gennevilliers, France) at the Centre for Microscopy, Characterisation and Analysis (The University of Western Australia). A 16-keV Cs+ primary ion beam of ∼0.75 pA (D1=3) focused to approximately 70 nm was rastered over the 25 μm^2^ sample area (256×256 pixel), with a dwell time of 60ms/pixel. Before analysis, each area was pre-implanted with a ∼3×10^16^ ions per cm^2^. Detectors (electron multipliers) were positioned to simultaneously measure negative secondary ions (^12^C^14^N, ^31^P^16^O_2_, ^34^S, ^12^C_2_). Mass resolving power was optimised using Entrance slit 3 (20µm), aperture slit 2 (200µm) and energy slit 1(∼10% yield reduction) and calculated as being ∼9000 (^12^C^14^N detector) according to Cameca’s MRP definition – sufficient to resolve all ion species of interest. Based on the secondary ion ^12^C^14^N count map, two regions of interest (ROI) were defined by manual drawing (parasite cell) and thresholding (host chromosomes) with the look@nanosims software (Polerecky et al., 2012). Ion counts (normalized by scans and pixels number) and ratios (^12^C^14^N/^12^C_2_, ^31^P^16^O_2_/^12^C_2_, ^34^S/^12^C_2_) were calculated for each ROI (Table S3). Ratio analyses do not provide absolute quantification of nitrogen (N), phosphorous (P) and sulfur (S) concentration but a comparison of the relative content of these elements between ROIs (host chromosomes vs parasite cell). In total, 131 host chromosomes were measured from 27 infected microalgal cells and 22 parasite cells.

### Transcriptomics analyses

#### Curating of enzymes involved in the energy metabolic pathways and sugar transport of the parasite *Amoebophyra* (syndiniales)

Reference proteins of interests were downloaded from the UniProtKB (https://www.uniprot.org) and VEuPathDB (https://veupathdb.org/veupathdb/app) databases (last access October 2021). Reference sequences for Sugars Will Eventually be Exported Transporters (SWEET) were obtained from a previous study (Jia et al., 2017). These reference sequences were used as BLAST queries to identify homologues in the *Amoebophrya* genome (available here: http://application.sb-roscoff.fr/blast/hapar/download.html) and the identity of positive hits was confirmed by (1) reverse-BLAST to the UniProtKB database (https://www.uniprot.org/blast/; last access October 2021); (2) sequence search in InterPro (http://www.ebi.ac.uk/interpro/; last access October 2021); (3) domain search with Pfam 34.0 (http://pfam.xfam.org/; last access October 2021); (4) phylogenetic analysis as described in (Kayal et al., 2020). Briefly, homologous sequences were downloaded from public databases, aligned with *Amoebophrya* sequences using mafft v. 7.407 (Katoh et al., 2019), and the alignments were filtered with Gblocks v. 0.91b (Castresana, 2000). Single gene phylogenetic trees were reconstructed for each alignment using RAxML v. 8.2.12 (Stamatakis, 2014) and the tree visually inspected using FigTree v1.4.4 (http://tree.bio.ed.ac.uk/software/figtree/). SWEET-like phylogeny was limited to Myzozoa (Apicomplexa + Dinoflagellata) given the very divergent sequences found in eukaryotes. For each gene, the presence of transmembrane domains was evaluated using the TMHMM Server v. 2.0 (http://www.cbs.dtu.dk/services/TMHMM/). Subcellular location and signal peptides were identified suing TargetP v. 2.0 (https://services.healthtech.dtu.dk/service.php?TargetP-2.0) and SignalP v. 5.0 (https://services.healthtech.dtu.dk/service.php?SignalP-5.0), respectively. For sugar transporters, we assumed positive identification when the number of TMs were similar to those of reference homologues, i.e. 7 for SWEET and 12 for hexose transporters and the phylogenetic position of the *Amoebophrya* gene fell within the *myzozoan* clade (Alveolata excluding ciliates).

#### Gene expression analysis

We used the DESeq2 differential expression analysis tool from the Trinity v2 package (Haas et al., 2013) to monitor the abundance of the genes of interest in the metatranscriptome (combining the transcriptomes of the host and parasite) produced by previous studies (Farhat et al., 2018; Kayal et al., 2020) throughout the infection and in dinospores. In short, filtered RNA-seq reads for each replicate (infected cells were sampled in triplicates every 6 h during a 36 h-long infection cycle) were separately mapped with the Bowtie2 aligner module of Trinity, and gene expression matrices were computed using the RSEM method (Li and Dewey, 2011). Not cross-sample normalized transcript per million (TPM) values were calculated for each species, and each time step separately. Only time steps corresponding to an average of 3x coverage of the transcriptome of the parasite were retained for interpretation in this study, namely two replicates at 18h (A and B) and all replicates for the following time steps (24, 30, 36 and dinospore), based on overall host and parasite remapping values (Table S4). For each gene, the time step where expression was maximum was estimated as the percentage of the average of replicates for each time step divided by the maximum average expression over the whole life cycle. Heatmaps of gene expression values were created using Heatmapper (http://www.heatmapper.ca/).

#### Identification of host transcripts

Known proteins (starch synthases, proteins of the FASII fatty acid biosynthesis pathway) of the dinoflagellate *Scrippsiella trochoidea*, closely related to the host (*Scrippsiella acuminata*), were used as queries against the un-infected host transcriptome. The respective best hit (or >1 hit, if several isoforms of one transcript were identified) were added to existing phylogenies for the corresponding protein (Hehenberger et al., 2016, 2019) to confirm the identity of the retrieved host transcripts. For the proteins involved in starch degradation, acetyl-CoA carboxylase (ACC) and the proteins involved in lipid droplet biosynthesis, new phylogenies were generated, using annotated proteins from *Arabidopsis thaliana* and/or *Homo sapiens* as queries (downloaded from https://www.uniprot.org; last access October 2021). The queries were used in a BLASTP search against a comprehensive custom database containing representatives from all major eukaryotic groups (except excavates) and RefSeq data from all bacterial phyla at NCBI (last accessed December 2017). The database was subjected to CD-HIT (https://pubmed.ncbi.nlm.nih.gov/23060610/) with a similarity threshold of 85% to reduce redundant sequences and paralogs. The search results of the BLASTP step were parsed for hits with an e-value threshold ≤1e-25 and a query coverage of ≥ 50% to reduce the possibility of paralogs and short sequences. The number of bacterial hits was restrained to 20 hits per phylum (for FCB group, most classes of Proteobacteria, PVC group, Spirochaetes, Actinobacteria, Cyanobacteria (unranked) and Firmicutes) or 10 per phylum (remaining bacterial phyla) as defined by NCBI taxonomy. Parsed hits were aligned with MAFFT v. 7.480, using the --auto option, poorly aligned regions were eliminated using trimAl v. 1.2 (https://pubmed.ncbi.nlm.nih.gov/19505945/) with a gap threshold of 80% and Maximum likelihood tree reconstructions were performed with FastTree v. 2.1.7 using the default options. The resulting phylogenies were inspected in FigTree v1.4.4, and recovered *S. trochoidea* hits were, as described above, used to identify homologs in the host transcriptome. Tree reconstruction and visual inspection were repeated including the host candidates. Complex, unresolved phylogenies (DGAT, ACAT) were further investigated by first manually inspecting the initial phylogenies and underlying alignments to remove contaminating, divergent and/or low-quality sequences. The cleaned, unaligned sequences were then subjected to filtering with PREQUAL using the default options (https://pubmed.ncbi.nlm.nih.gov/29868763/) to remove non homologous residues introduced by poor-quality sequences, followed by alignment with MAFFT G-INS-i using the VSM option (--unalignlevel 0.6) to control over-alignment (Katoh et al., 2019). The alignments were subjected to Divvier (Ali et al., 2019) using the -divvygap and the -mincol 4 option to improve homology inference before removing ambiguously aligned sites with trimAl (-gt 0.01). Final trees were calculated with IQ-TREE v. 1.6.5 (Nguyen et al., 2015), using the -mset option to restrict model selection to LG for ModelFinder (Kalyaanamoorthy et al., 2017), while branch support was assessed with 1000 ultrafast bootstrap replicates (Hoang et al., 2018). In addition, all putative DGAT and ACAT candidates in the host were submitted to InterProScan at https://www.ebi.ac.uk/interpro/ to confirm their identity (Blum et al., 2021).

## Acknowledgements

J.D. was supported by CNRS and ATIP-Avenir program. This project has received financial support from the CNRS through the MITI interdisciplinary programs. This project also received funding from the LabEx GRAL (ANR-10-LABX-49-01), financed within the University Grenoble Alpes graduate school (Ecoles Universitaires de Recherche) CBH-EUR-GS (ANR-17-EURE-0003). This project was also supported by the European Union’s Horizon 2020 research and innovation programme CORBEL under the grant agreement No 654248. This work used the platforms of the Grenoble Instruct-ERIC centre (ISBG; UAR 3518 CNRS-CEA-UGA-EMBL) within the Grenoble Partnership for Structural Biology (PSB), supported by FRISBI (ANR-10-INBS-05-02) and GRAL, financed within the University Grenoble Alpes graduate school (Ecoles Universitaires de Recherche) CBH-EUR-GS (ANR-17-EURE-0003). We thank Guy Schoehn and Christine Moriscot, and the electron microscope facility, which is supported by the Rhône-Alpes Region, the Fondation Recherche Medicale (FRM), the fonds FEDER, the Centre National de la Recherche Scientifique (CNRS), the CEA, the University of Grenoble, EMBL, and the GIS-Infrastructures en Biologie Sante et Agronomie (IBISA). E.K. acknowledges the Institut Français de Bioinformatique (ANR-11-INBS-0013) and the Roscoff Bioinformatics platform ABiMS (http://abims.sb-roscoff.fr) for providing computing and storage resources. E.K was supported by the French Government via the National Research Agency investment expenditure program IDEALG (ANR-10-BTBR-04). We acknowledge use of the Microscopy Australia facilities at the Centre for Microscopy, Characterisation & Analysis, The University of Western Australia, a facility funded by the University, State and Commonwealth Governments.

## Notes

### Competing Interest Statement

The authors have declared no competing interest.

## References

Alacid, E., Reñé, A., and Garcés, E. (2015). New Insights into the Parasitoid Parvilucifera sinerae Life Cycle: The Development and Kinetics of Infection of a Bloom-forming Dinoflagellate Host. Protist 166, 677–699.

Ali, R.H., Bogusz, M., Whelan, S., and Tamura, K. (2019). Identifying Clusters of High Confidence Homologies in Multiple Sequence Alignments. Mol. Biol. Evol. 36, 2340–2351.

Amiar, S., Katris, N.J., Berry, L., Mcfadden, G.I., Botte, C.Y., Amiar, S., Katris, N.J., Berry, L., Dass, S., Duley, S., et al. (2020). Division and Adaptation to Host Environment of Apicomplexan Parasites Depend on Apicoplast Lipid Metabolic Plasticity and Host Organelle Remodeling Article Division and Adaptation to Host Environment of Apicomplexan Parasites Depend on Apicoplast Lipid Me. 3778–3792.

Blum, M., Chang, H.Y., Chuguransky, S., Grego, T., Kandasaamy, S., Mitchell, A., Nuka, G., Paysan-Lafosse, T., Qureshi, M., Raj, S., et al. (2021). The InterPro protein families and domains database: 20 years on. Nucleic Acids Res. 49, D344–D354.

Blume, M., Rodriguez-Contreras, D., Landfear, S., Fleige, T., Soldati-Favre, D., Lucius, R., and Gupta, N. (2009). Host-derived glucose and its transporter in the obligate intracellular pathogen Toxoplasma gondii are dispensable by glutaminolysis. Proc. Natl. Acad. Sci. U. S. A. 106, 12998–13003.

Cachon, J. (1964). Contribution à l’étude des péridiniens parasites. Cytologie, cycles évolutifs. Ann. Sci. Nat. Zool 6, 1–158.

Caffaro, C.E., and Boothroyd, J.C. (2011). Evidence for host cells as the major contributor of lipids in the intravacuolar network of toxoplasma-infected cells. Eukaryot. Cell 10, 1095–1099.

Cai, R., Kayal, E., Alves-de-Souza, C., Bigeard, E., Corre, E., Jeanthon, C., Marie, D., Porcel, B.M., Siano, R., Szymczak, J., et al. (2020). Cryptic species in the parasitic Amoebophrya species complex revealed by a polyphasic approach. Sci. Rep. 10, 1–11.

Castresana, J. (2000). Selection of Conserved Blocks from Multiple Alignments for Their Use in Phylogenetic Analysis. Mol. Biol. Evol. 17, 540–552.

Chen, L.Q., Qu, X.Q., Hou, B.H., Sosso, D., Osorio, S., Fernie, A.R., and Frommer, W.B. (2012). Sucrose efflux mediated by SWEET proteins as a key step for phloem transport. Science (80-.). 335, 207–211.

Coats, D.W. (1999). Parasitic life styles of marine dinoflagellates. J. Eukaryot. Microbiol. 46, 402–409.

Coats, D.W., and Park, M.G. (2002). Parasitism of photosynthetic dinoflagellates by three strains of Amoebophrya (Dinophyta): Parasite survival, infectivity, generation time, and host specificity. J. Phycol. 38, 520–528.

Cox, D., Lee, D.J., Dale, B.M., Calafat, J., and Greenberg, S. (2000). A Rab11-containing rapidly recycling compartment in macrophages that promotes phagocytosis. Proc. Natl. Acad. Sci. U. S. A. 97, 680–685.

Decelle, J., Stryhanyuk, H., Gallet, B., Veronesi, G., Schmidt, M., Balzano, S., Marro, S., Uwizeye, C., Jouneau, P.H., Lupette, J., et al. (2019). Algal Remodeling in a Ubiquitous Planktonic Photosymbiosis. Curr. Biol. 29, 968-978.e4.

Van Dooren, G.G., Marti, M., Tonkin, C.J., Stimmler, L.M., Cowman, A.F., and McFadden, G.I. (2005). Development of the endoplasmic reticulum, mitochondrion and apicoplast during the asexual life cycle of Plasmodium falciparum. Mol. Microbiol. 57, 405–419.

Elliott, D.A., McIntosh, M.T., Hosgood, H.D., Chen, S., Zhang, G., Baevova, P., and Joiner, K.A. (2008). Four distinct pathways of hemoglobin uptake in the malaria parasite Plasmodium falciparum. Proc. Natl. Acad. Sci. U. S. A. 105, 2463–2468.

Evers, F., Cabrera-Orefice, A., Elurbe, D.M., Keate Lindert, M., Boltryk, S.D., Voss, T.S., Huynen, M.A., Brandt, U., and Kooij, T.W.A. (2021). Composition and stage dynamics of mitochondrial complexes in Plasmodium falciparum. Nat. Commun. 12, 1–17.

Farhat, S., Florent, I., Noel, B., Kayal, E., Da Silva, C., Bigeard, E., Alberti, A., Labadie, K., Corre, E., Aury, J.M., et al. (2018). Comparative time-scale gene expression analysis highlights the infection processes of two amoebophrya strains. Front. Microbiol. 9, 1–19.

Farhat, S., Le, P., Kayal, E., Noel, B., Bigeard, E., Corre, E., Maumus, F., Florent, I., Alberti, A., Aury, J.M., et al. (2021). Rapid protein evolution, organellar reductions, and invasive intronic elements in the marine aerobic parasite dinoflagellate Amoebophrya spp. BMC Biol. 19, 1–21.

Flegontov, P., Michálek, J., Janouškovec, J., Lai, D.H., Jirků, M., Hajdušková, E., Tom čala, A., Otto, T.D., Keeling, P.J., Pain, A., et al. (2015). Divergent mitochondrial respiratory chains in phototrophic relatives of apicomplexan parasites. Mol. Biol. Evol. 32, 1115–1131.

Fox, B.A., Guevara, R.B., Rommereim, L.M., Falla, A., Bellini, V., Pètre, G., Rak, C., Cantillana, V., Dubremetz, J.-F., Cesbron-Delauw, M.-F., et al. (2019). Toxoplasma gondii Parasitophorous Vacuole Membrane-Associated Dense Granule Proteins Orchestrate Chronic Infection and GRA12 Underpins Resistance to Host Gamma Interferon. MBio 10.

Freeman Rosenzweig, E.S., Xu, B., Kuhn Cuellar, L., Martinez-Sanchez, A., Schaffer, M., Strauss, M., Cartwright, H.N., Ronceray, P., Plitzko, J.M., Förster, F., et al. (2017). The Eukaryotic CO2-Concentrating Organelle Is Liquid-like and Exhibits Dynamic Reorganization. Cell 171, 148-162.e19.

Gomes, A.F., Magalhães, K.G., Rodrigues, R.M., de Carvalho, L., Molinaro, R., Bozza, P.T., and Barbosa, H.S. (2014). Toxoplasma gondii-skeletal muscle cells interaction increases lipid droplet biogenesis and positively modulates the production of IL-12, IFN-g and PGE2. Parasit. Vectors 7, 47.

Guillou, L., Viprey, M., Chambouvet, A., Welsh, R.M., Kirkham, A.R., Massana, R., Scanlan, D.J., and Worden, A.Z. (2008). Widespread occurrence and genetic diversity of marine parasitoids belonging to Syndiniales (Alveolata). 10, 3349–3365.

Haas, B.J., Papanicolaou, A., Yassour, M., Grabherr, M., Blood, P.D., Bowden, J., Couger, M.B., Eccles, D., Li, B., Lieber, M., et al. (2013). De novo transcript sequence reconstruction from RNA-seq using the Trinity platform for reference generation and analysis. Nat. Protoc. 8, 1494–1512.

Harris, E., Yoshida, K., Cardelli, J., and Bush, J. (2001). Rab11-like GTPase associates with and regulates the structure and function of the contractile vacuole system in Dictyostelium. J. Cell Sci. 114, 3035–3045.

Hehenberger, E., Burki, F., Kolisko, M., and Keeling, P.J. (2016). Functional Relationship between a Dinoflagellate Host and Its Diatom Endosymbiont. Mol. Biol. Evol. 33, 2376–2390.

Hehenberger, E., Gast, R.J., and Keeling, P.J. (2019). A kleptoplastidic dinoflagellate and the ipping point between transient and fully integrated plastid endosymbiosis. Proc. Natl. Acad. Sci. U. S. A. 116, 17934–17942.

Hennies, J., Lleti, J.M.S., Schieber, N.L., Templin, R.M., Steyer, A.M., and Schwab, Y. (2020). AMST: Alignment to Median Smoothed Template for Focused Ion Beam Scanning Electron Microscopy Image Stacks. Sci. Rep. 10, 1–10.

Hoang, D.T., Chernomor, O., von Haeseler, A., Minh, B.Q., and Vinh, L.S. (2018). UFBoot2: Improving the Ultrafast Bootstrap Approximation. Molecular biology and evolution. Mol. Biol. Evol. 35, 518–522.

Hu, X., Binns, D., and Reese, M.L. (2017). The coccidian parasites Toxoplasma and Neospora dysregulate mammalian lipid droplet biogenesis. J. Biol. Chem. 292, 11009–11020.

Jacot, D., Waller, R.F., Soldati-Favre, D., MacPherson, D.A., and MacRae, J.I. (2016). Apicomplexan Energy Metabolism: Carbon Source Promiscuity and the Quiescence Hyperbole. Trends Parasitol. 32, 56–70.

Jephcott, T.G., Alves-de-Souza, C., Gleason, F.H., van Ogtrop, F.F., Sime-Ngando, T., Karpov, S.A., and Guillou, L. (2016). Ecological impacts of parasitic chytrids, syndiniales and perkinsids on populations of marine photosynthetic dinoflagellates. Fungal Ecol. 19, 47–58.

Jia, B., Zhu, X.F., Pu, Z.J., Duan, Y.X., Hao, L.J., Zhang, J., Chen, L.Q., Jeon, C.O., and Xuan, Y.H. (2017). Integrative view of the diversity and evolution of SWEET and semiSWEET sugar transporters. Front. Plant Sci. 8, 1–18.

John, U., Lu, Y., Wohlrab, S., Groth, M., Janouškovec, J., Kohli, G.S., Mark, F.C., Bickmeyer, U., Farhat, S., Felder, M., et al. (2019). An aerobic eukaryotic parasite with functional mitochondria that likely lacks a mitochondrial genome. Sci. Adv. 5, eaav1110.

Kalyaanamoorthy, S., Minh, B.Q., Wong, T.K.F., von Haeseler, A., and Jermiin, L.S. (2017). ModelFinder: fast model selection for accurate phylogenetic estimates. Nat. Methods 14, 587–589.

Katoh, K., Rozewicki, J., and Yamada, K.D. (2019). MAFFT online service: multiple sequence alignment, interactive sequence choice and visualization. Brief. Bioinform. 20, 1160–1166.

Kayal, E., Alves-de-Souza, C., Farhat, S., Velo-Suarez, L., Monjol, J., Szymczak, J., Bigeard, E., Marie, D., Noel, B., Porcel, B.M., et al. (2020). Dinoflagellate Host Chloroplasts and Mitochondria Remain Functional During Amoebophrya Infection. Front. Microbiol. 11, 1–11.

Kikinis, R., Pieper, S.D., and Vosburgh, K.G. (2014). 3D Slicer: A Platform for Subject-Specific Image Analysis, Visualization, and Clinical Support. In Intraoperative Imaging and Image-Guided Therapy, (New York, NY: Springer New York), pp. 277–289.

Krungkrai, J. (2004). The multiple roles of the mitochondrion of the malarial parasite. 511–524.

Latorraca, N.R., Fastman, N.M., Venkatakrishnan, A.J., Frommer, W.B., Dror, R.O., and Feng, L. (2017). Mechanism of Substrate Translocation in an Alternating Access Transporter. Cell 169, 96-107.e12.

Lee, J.W. (2020). Protonic Capacitor: Elucidating the biological significance of mitochondrial cristae formation. Sci. Rep. 10, 1–14.

Li, B., and Dewey, C.N. (2011). RSEM: accurate transcript quantification from RNA-Seq data with or without a reference genome. BMC Bioinformatics 12, 323.

Lima-Mendez, G., Faust, K., Henry, N., Decelle, J., Colin, S., Carcillo, F., Chaffron, S., Ignacio-Espinosa, J.C., Roux, S., Vincent, F., et al. (2015). Determinants of community structure in the global plankton interactome. Science 348, 1262073_1-1262073_9.

Lopez, J., Bittame, A., Massera, C., Vasseur, V., Effantin, G., Valat, A., Buaillon, C., Allart, S., Fox, B.A.A., Rommereim, L.M.M., et al. (2015). Intravacuolar Membranes Regulate CD8 T Cell Recognition of Membrane-Bound Toxoplasma gondii Protective Antigen. Cell Rep. 13, 2273–2286.

Lowe, D.G. (2004). Distinctive Image Features from Scale-Invariant Keypoints. Int. J. Comput. Vis. 60, 91–110.

Marshansky, V., Rubinstein, J.L., and Grüber, G. (2014). Eukaryotic V-ATPase: Novel structural findings and functional insights. Biochim. Biophys. Acta - Bioenerg. 1837, 857–879.

McFadden, G.I., Reith, M.E., Munholland, J., and Lang-Unnasch, N. (1996). Plastid in human parasites. Nature 381, 482.

Miller, J.J., Delwiche, C.F., and Coats, D.W. (2012). Ultrastructure of Amoebophrya sp. and its Changes during the Course of Infection. Ann. Anat. 163, 720–745.

Mühleip, A., Kock Flygaard, R., Ovciarikova, J., Lacombe, A., Fernandes, P., Sheiner, L., and Amunts, A. (2021). ATP synthase hexamer assemblies shape cristae of Toxoplasma mitochondria. Nat. Commun. 12.

Nguyen, L.T., Schmidt, H.A., Von Haeseler, A., and Minh, B.Q. (2015). IQ-TREE: A fast and effective stochastic algorithm for estimating maximum-likelihood phylogenies. Mol. Biol. Evol. 32, 268–274.

Nishi, M., Hu, K., Murray, J.M., and Roos, D.S. (2008). Organellar dynamics during the cell cycle of Toxoplasma gondii. J. Cell Sci. 121, 1559–1568.

Nolan, S.J., Romano, J.D., and Coppens, I. (2017). Host lipid droplets: An important source of lipids salvaged by the intracellular parasite Toxoplasma gondii. PLoS Pathog. 13, e1006362.

Painter, H.J., Morrisey, J.M., Mather, M.W., and Vaidya, A.B. (2007). Specific role of mitochondrial electron transport in. 446.

Pánek, T., Eliáš, M., Vancová, M., Lukeš, J., and Hashimi, H. (2020). Returning to the Fold for Lessons in Mitochondrial Crista Diversity and Evolution. Curr. Biol. 30, R575–R588.

Pinchuk, G.E., Ammons, C., Culley, D.E., Li, S.M.W., McLean, J.S., Romine, M.F., Nealson, K.H., Fredrickson, J.K., and Beliaev, A.S. (2008). Utilization of DNA as a sole source of phosphorus, carbon, and energy by Shewanella spp.: Ecological and physiological implications for dissimilatory metal reduction. Appl. Environ. Microbiol. 74, 1198–1208.

Polerecky, L., Adam, B., Milucka, J., Musat, N., Vagner, T., and Kuypers, M.M.M. (2012). Look@NanoSIMS - a tool for the analysis of nanoSIMS data in environmental microbiology. Environ. Microbiol. 14, 1009–1023.

Prapunwattana, P., and Krungkrai, S.R. (1999). Ultrastructure and function of mitochondria in gametocytic stage of.

Pszenny, V., Ehrenman, K., Romano, J.D., Kennard, A., Schultz, A., Roos, D.S., Grigg, M.E., Carruthers, V.B., and Coppens, I. (2016). A lipolytic lecithin:Cholesterol acyltransferase secreted by toxoplasma facilitates parasite replication and egress. J. Biol. Chem. 291, 3725–3746.

Qureshi, A.A., Suades, A., Matsuoka, R., Brock, J., McComas, S.E., Nji, E., Orellana, L., Claesson, M., Delemotte, L., and Drew, D. (2020). The molecular basis for sugar import in malaria parasites. Nature 578, 321–325.

Salcedo-Sora, J.E., Caamano-Gutierrez, E., Ward, S.A., and Biagini, G.A. (2014). The proliferating cell hypothesis: A metabolic framework for Plasmodium growth and development. Trends Parasitol. 30, 170–175.

Siano, R., Alves-De-Souza, C., Foulon, E. M. Bendif, E., Simon, N., Guillou, L., and Not, F. (2011). Distribution and host diversity of Amoebophryidae parasites across oligotrophic waters of the Mediterranean Sea. Biogeosciences 8, 267–278.

Sibbald, S.J., and Archibald, J.M. (2020). Genomic Insights into Plastid Evolution. Genome Biol. Evol. 12, 978–990.

Stamatakis, A. (2014). RAxML version 8: a tool for phylogenetic analysis and post-analysis of large phylogenies. Bioinformatics 30, 1312–1313.

Stephan, T., Brüser, C., Deckers, M., Steyer, A.M., Balzarotti, F., Barbot, M., Behr, T.S., Heim, G., Hübner, W., Ilgen, P., et al. (2020). MICOS assembly controls mitochondrial inner membrane remodeling and crista junction redistribution to mediate cristae formation. EMBO J. 39, 1–24.

Uwizeye, C., Mars, M., Gallet, B., Chevalier, F., and Lekieffre, C. (2021a). Cytoklepty in the plankton □ : A host strategy to optimize the bioenergetic machinery of endosymbiotic algae. 118.

Uwizeye, C., Decelle, J., Jouneau, P., Flori, S., Gallet, B., Keck, J., Bo, D.D., Moriscot, C., Seydoux, C., Chevalier, F., et al. (2021b). Morphological bases of phytoplankton energy management and physiological responses unveiled by 3D subcellular imaging. Nat. Commun. 12, 1049.

de Vargas, C., Audic, S., Henry, N., Decelle, J., Mahe, F., Logares, R., Lara, E., Berney, C., Le Bescot, N., Probert, I., et al. (2015). Eukaryotic plankton diversity in the sunlit ocean. Science 348, 1261605.

Vines, J.H., and King, J.S. (2019). The endocytic pathways of Dictyostelium discoideum. Int. J. Dev. Biol. 63, 461–471.

Voleman, L., and Doležal, P. (2019). Mitochondrial dynamics in parasitic protists. PLoS Pathog. 15.

Wideman, J.G., and Muñoz-gómez, S.A. (2016). The evolution of ERMIONE in mitochondrial biogenesis and lipid homeostasis □: An evolutionary view from comparative cell biology ⍰. Biochim. Biophys. Acta 1861, 900–912.

Worden, A.Z., Follows, M.J., Giovannoni, S.J., Wilken, S., Zimmerman, A.E., and Keeling, P.J. (2015). Rethinking the marine carbon cycle: Factoring in the multifarious lifestyles of microbes. Science (80-.). 347, 1257594–1257594.

Zeeman, S.C., Kossmann, J., and Smith, A.M. (2010). Starch: Its metabolism, evolution, and biotechnological modification in plants. Annu. Rev. Plant Biol. 61, 209–234.

