## Supplementary figures for "Intracellular development and impact of a eukaryotic parasite on its zombified microalgal host in the marine plankton"

**Supplementary Tables and Figures**

**Table S1:** Morphometric data of the organelles and compartments of the host (the dinoflagellate *Scrippsiella acuminata*) and the parasite *Amoebophyra* obtained after FIB-SEM-based 3D reconstructions.

**Table S2:** Transcriptomics data showing the expression level of genes of the parasite *Ameobophyra* at different infection stages.

**Table S3:** NanoSIMS (Nanoscale Secondary Ion Mass Spectrometry) data with natural abundance of phosphorous (^31^P^16^O_2_/^12^C_2_), sulfur (^34^S/^12^C_2_) and nitrogen (^12^C^14^N/^12^C_2_) measured on resin sections of an infected culture of *Scrippsiella acuminata* with the parasite *Amoebophyra*.

**Table S4:** Transcriptomics data of the host (*Scrippsiella acuminata*) with expression of genes involved in starch and lipids metabolism.

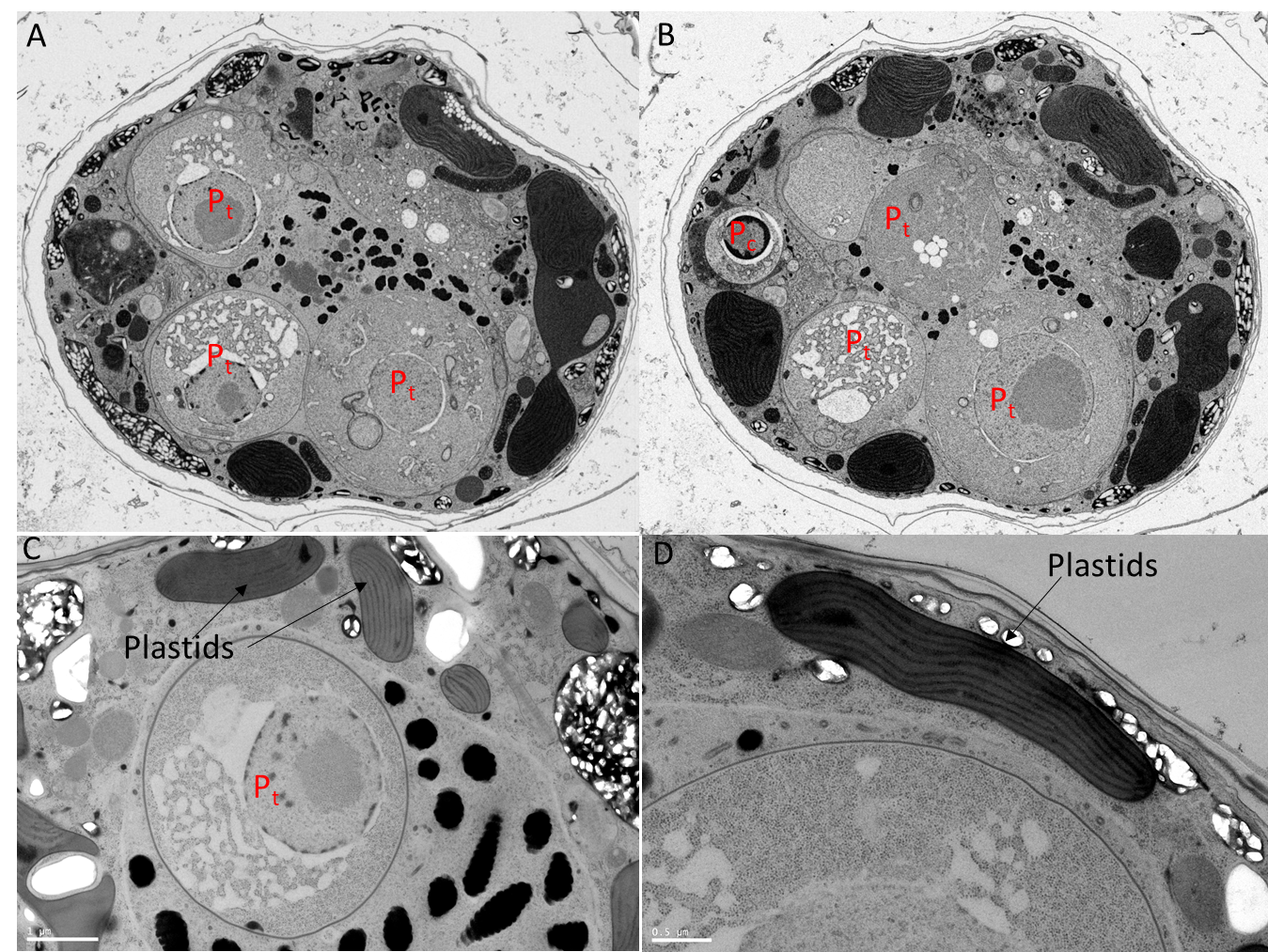

**Figure S1. Transmission Electron Microscopy (TEM) micrographs of the microalga *Scrippsiella acuminata* (dinoflagellate) infected by the parasite *Ameobophyra* sp. (syndiniales). A-B)** Multiple parasites at different developmental stages can infect simultaneously the host cell (P_c_: cytoplasmic stage and P_t_: trophont stages). **C-D)** During infection, the morphology of the host plastids remains intact, more particularly the arrangement of the thylakoids.

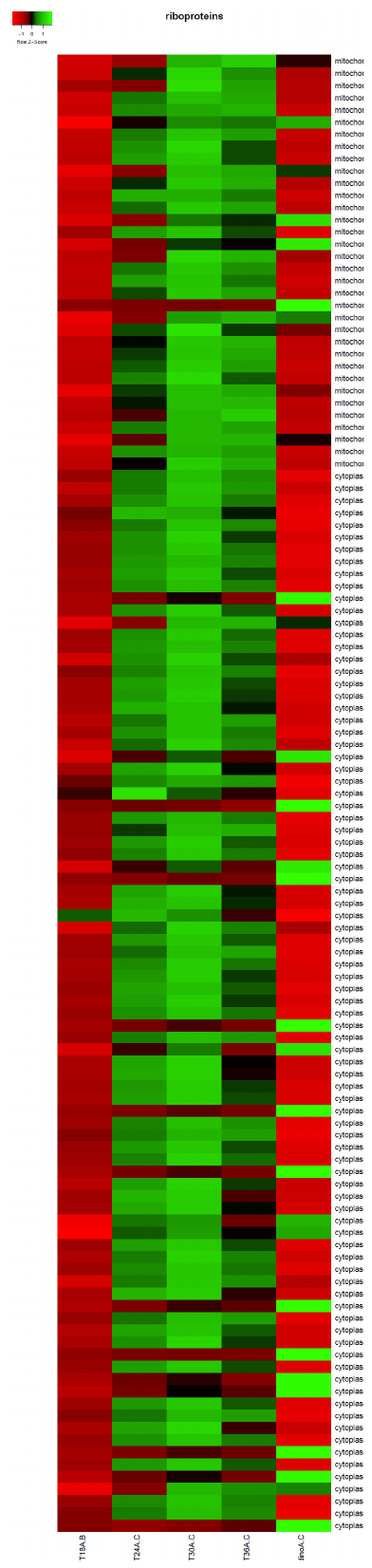

**Figure S2. Expression levels of the nuclear ribosomal genes of the parasite across different intracellular stages within its host and in dinospores.** Heatmap showing the expression level of nuclear ribosomal genes of the parasite during the infection (T18A-C, T24A-C, T30A-C and T36A-C) and the dinospore stage (dino).

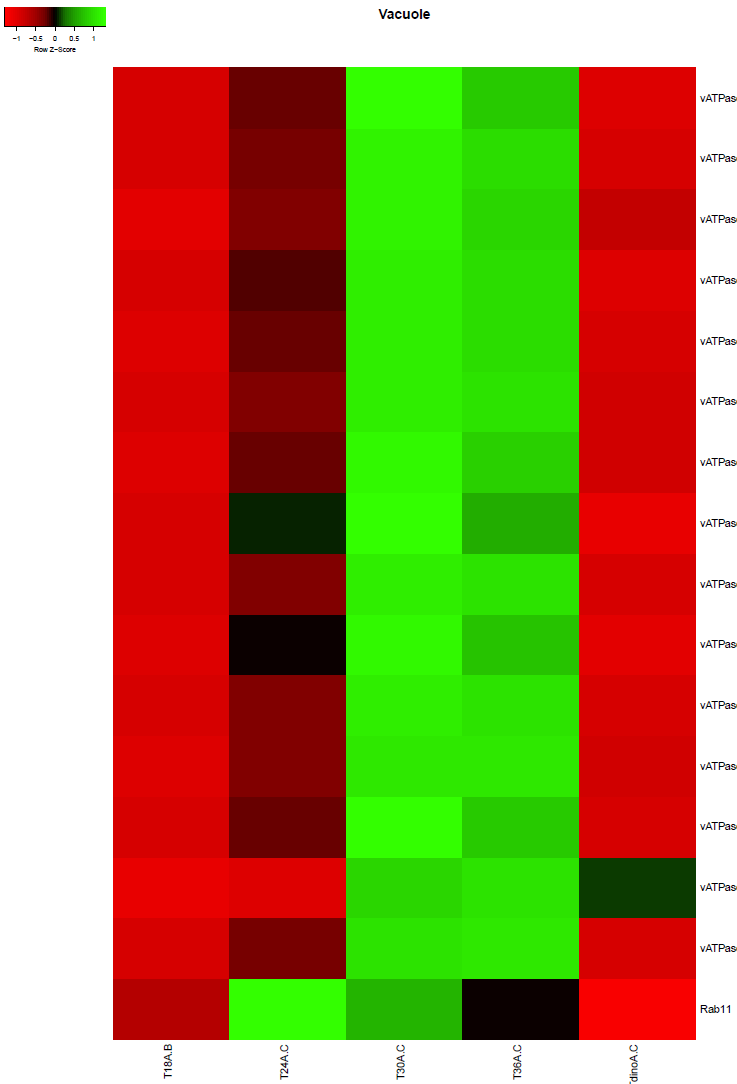

**Figure S3. Expression levels of the genes of the parasite potentially involved in the formation of phagotrophic vacuoles across different intracellular stages within its host and in dinospores.** Heatmap showing the expression level of subunits of the vacuolar H^+^-ATPase (V-ATPase) and the GTPase Ras-related protein Rab-11 at different intracellular stages (T18A-C, T24A-C, T30A-C and T36A-C) and in dinospores (dinoA-C).

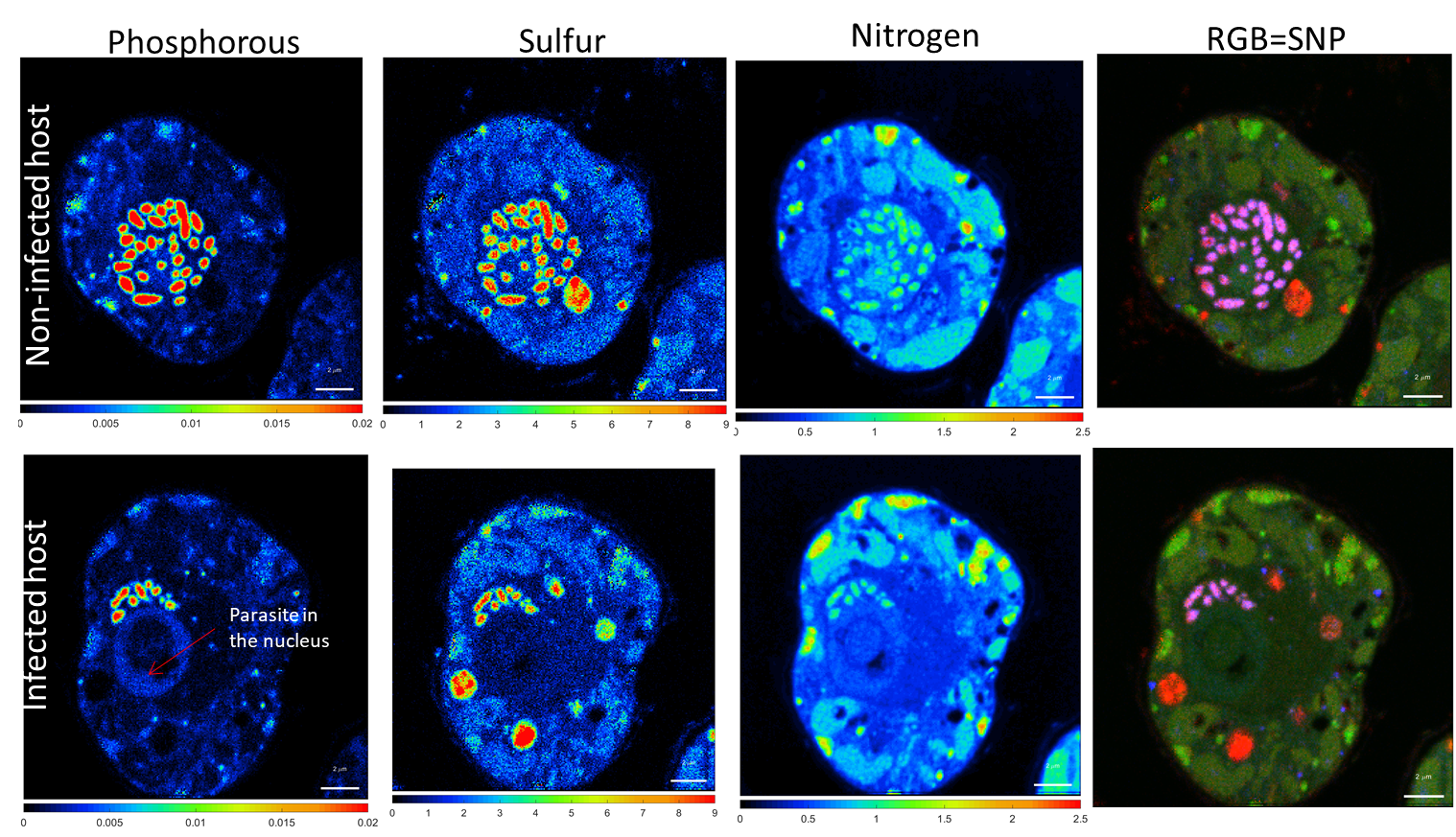

**Figure S4.** **Subcellular quantitative mapping of phosphorous, sulfur and nitrogen in non-infected hosts (the microalga *Scrippsiella acuminata*) (upper row) and infected hosts (lower row) with their intranuclear parasite unveiled by nanoSIMS (Nanoscale Secondary Ion Mass Spectrometry).** Subcellular distribution of phosphorous (^31^P^16^O_2_/^12^C_2_), nitrogen (^12^C^14^N/^12^C_2_), and sulfur (^34^S/^12^C_2_) and the overlapping of the three nutrients (RGB: red for sulfur, green for nitrogen and blue for phosphorous).The scale bar represents 2 µm.

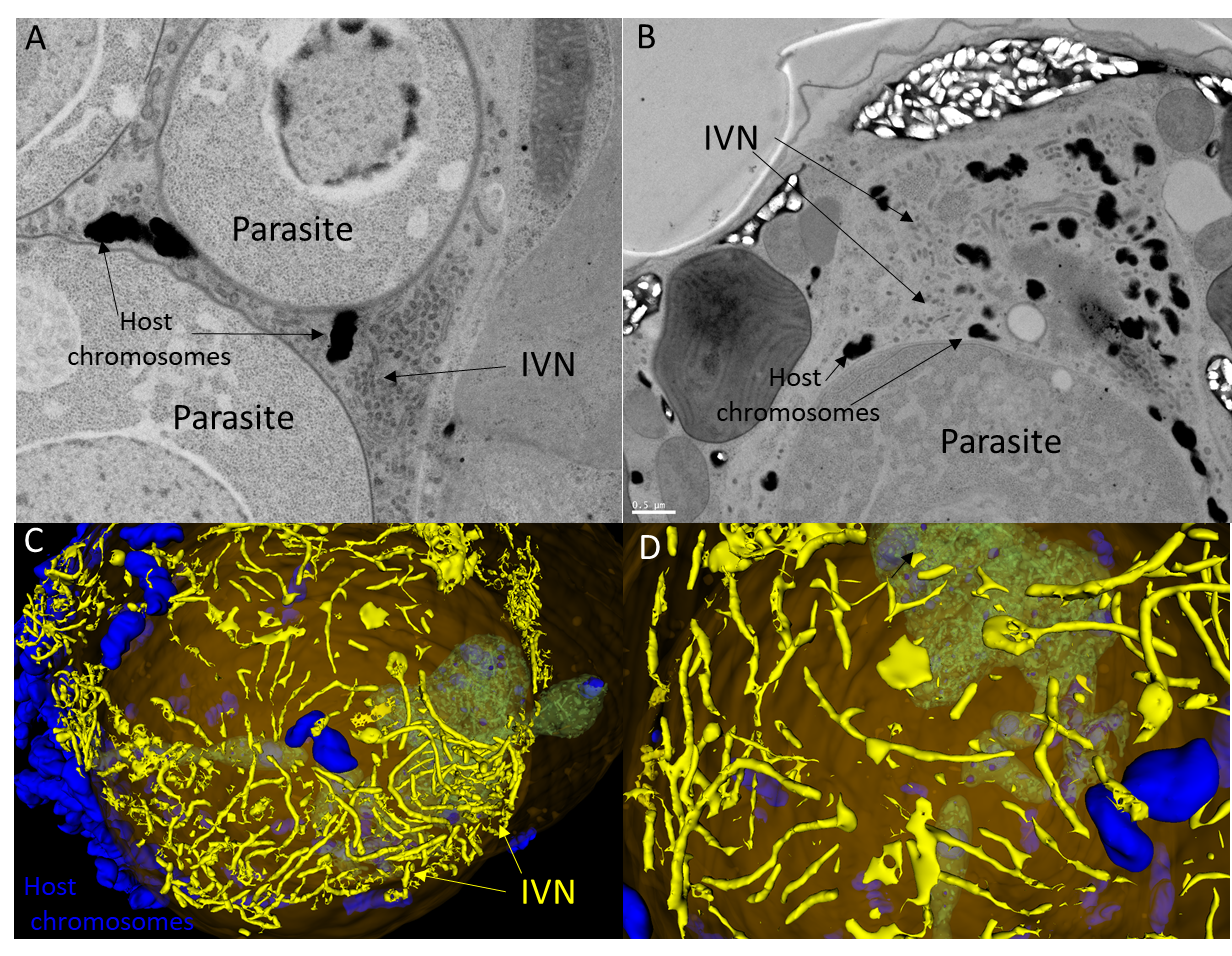

**Figure S5: Formation of a network of tubules in the host nucleus, which resembles the Intravacuolar Network (IVN) described in the human parasite *Toxoplasma.*** **A-B)** Micrographs of TEM (Transmission Electron Microscopy) showing that the IVN-like structures often surrounded and concentrated around the host chromosomes, more particularly at the trophont stages. **C-D)** 3D reconstructions of the IVN-like structures (yellow) and the host chromosomes (blue) after FIB-SEM (Focused-Ion Beam – Scanning Electron Microscopy).

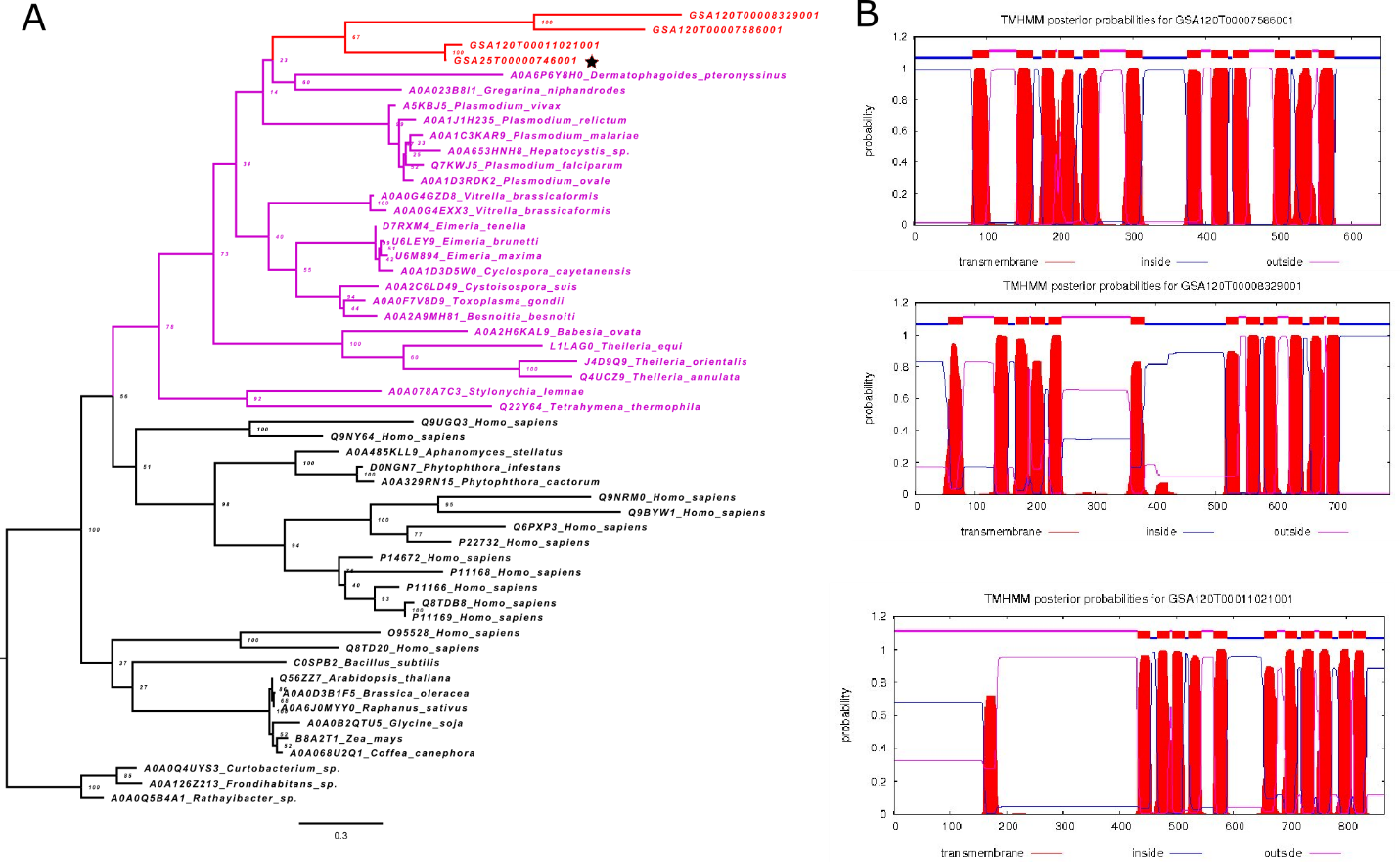

**Figure S6: Hexose transporters identified in the syndiniales parasite *Amoebophrya* sp. strains A120 and A25**. A) Maximum Likelihood phylogenetic analysis of hexose amino acid sequences using RAxML v.8 with the GTR+Γ model of sequence evolution and 100 bootstrap replicates, rooted by Bacteria. Alveolate and *Amoebophrya* sequences are highlighted in purple and red, respectively. A star represents the only hexose sequence identified in the *Amoebophrya* A25 strain not used in this study. The other three sequences represent the three hexose transporters HT1, HT2 and HT3 found in *Amoebophrya* A120. B) Predicted transmembrane helices of the three hexose genes identified in A120 using the TMHMM Server v. 2.0 (http://www.cbs.dtu.dk/services/TMHMM/).

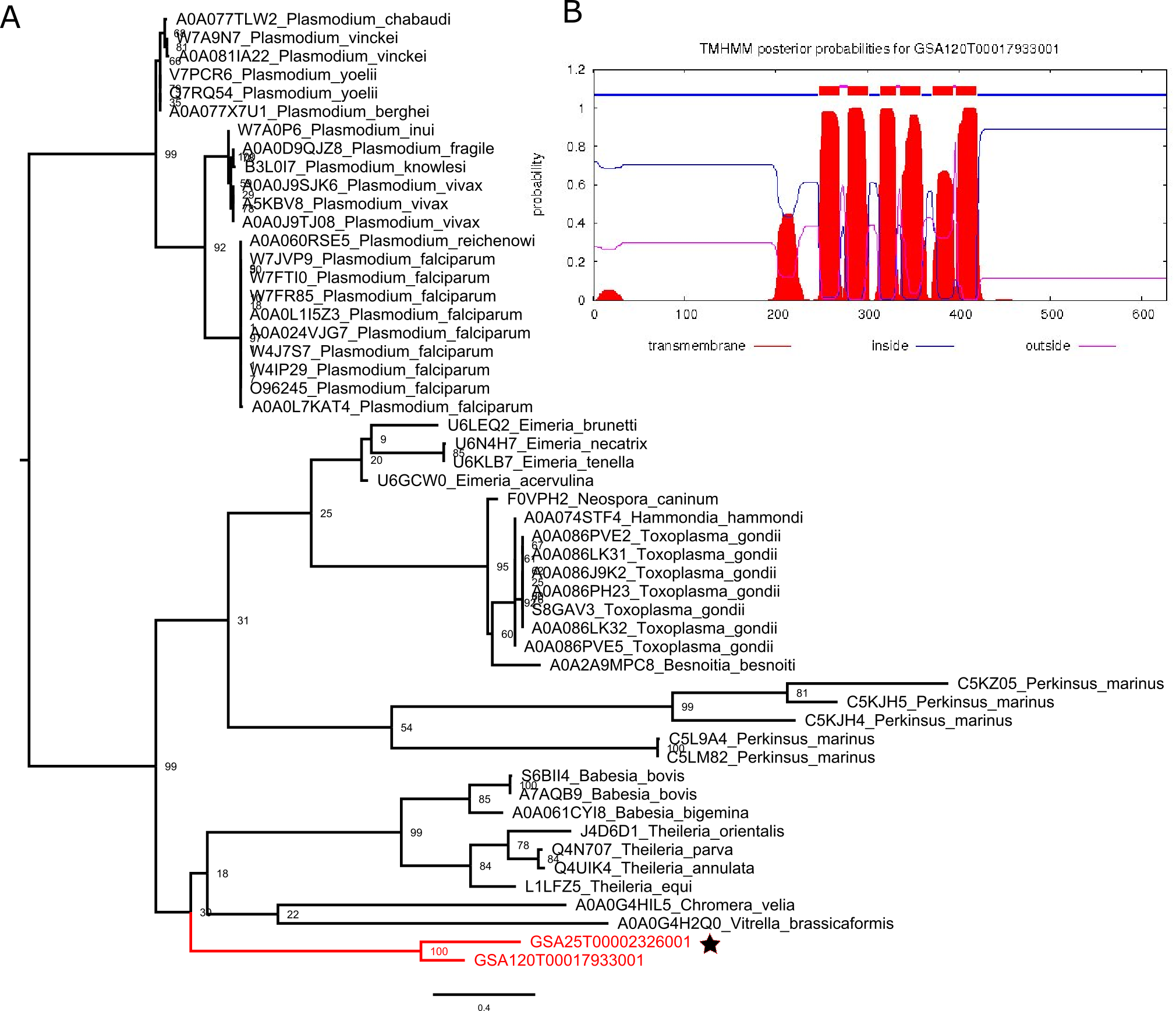

**Figure S7: SWEET transporters identified in the syndiniales parasite *Amoebophrya* sp. strains A120 and A25.** A) Maximum Likelihood phylogenetic analysis of SWEET amino acid sequences found in alveolates using RAxML v.8 with the GTR+Γ model of sequence evolution and 100 bootstrap replicates. *Amoebophrya* sequences are highlighted in red; a star denotes the SWEET sequence identified in the *Amoebophrya* A25 strain not used in this study. B) Predicted transmembrane helices of the SWEET gene identified in A120 using the TMHMM Server v. 2.0 (http://www.cbs.dtu.dk/services/TMHMM/).

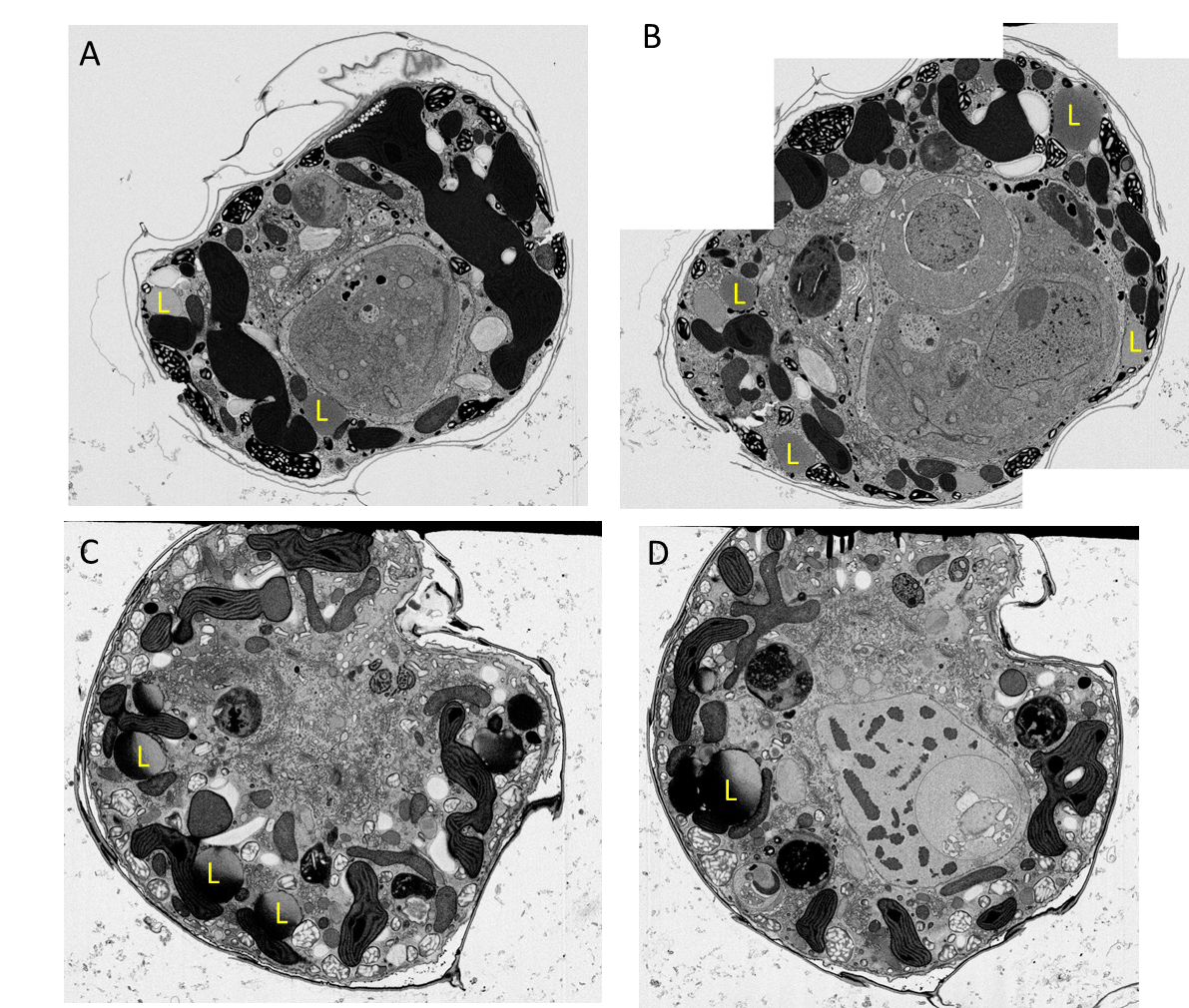

**Figure S8: Electron micrographs showing the formation of lipid droplets (L) in the host cell (the microalga *Scrippsiella acuminata*) during infection of the parasite *Amoebophyra* (Syndiniales).** Note that the lipid droplets (L) were in close association with plastids and mitochondrion of the host.

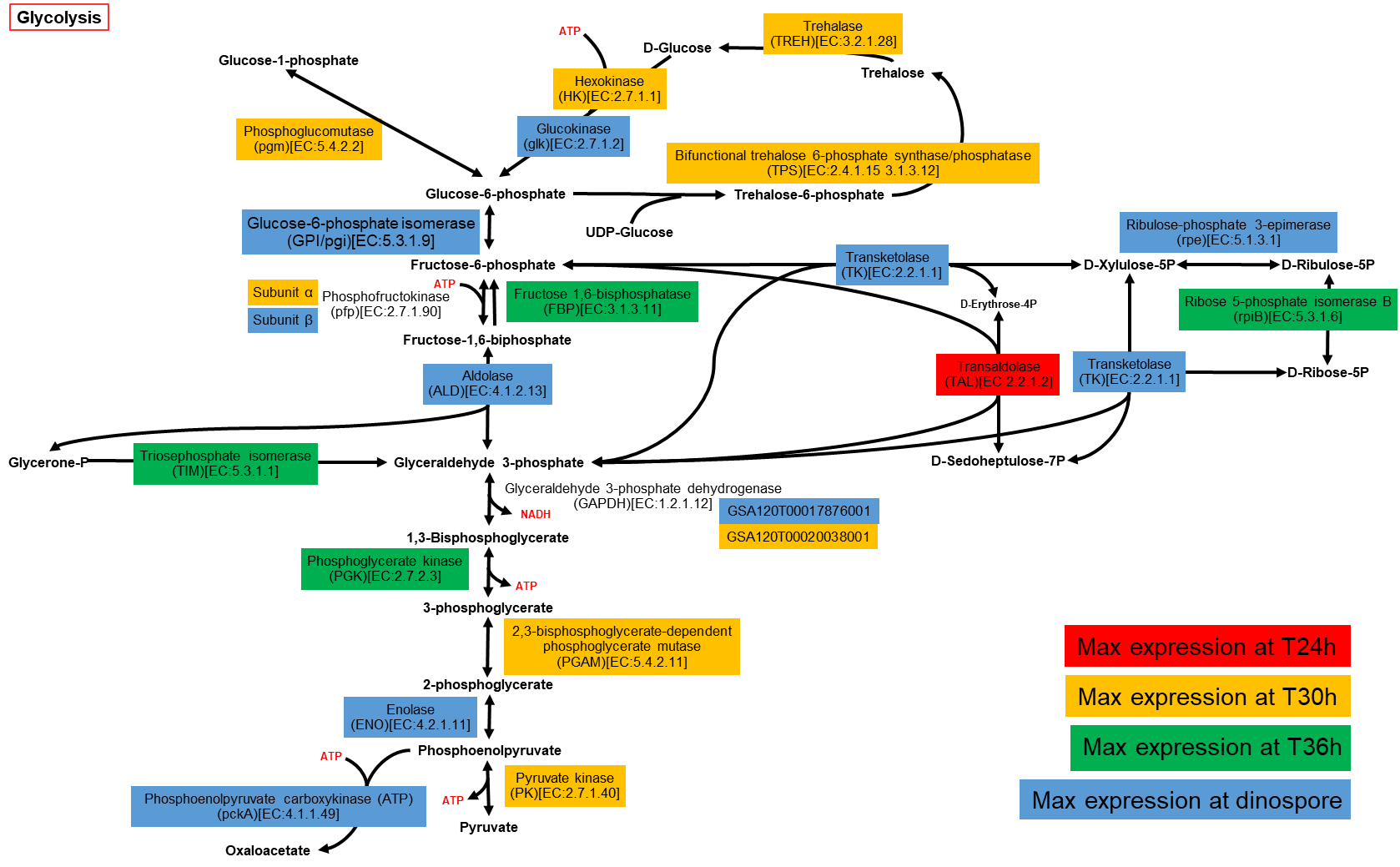

**Figure S9: Reconstruction of the glycolysis pathway in the syndiniales parasite *Amoebophrya* sp. strains A120 based on the genome.** Genes are colour-coded based on maximum expression estimated by averaging RNA-seq mapping for triplicates.

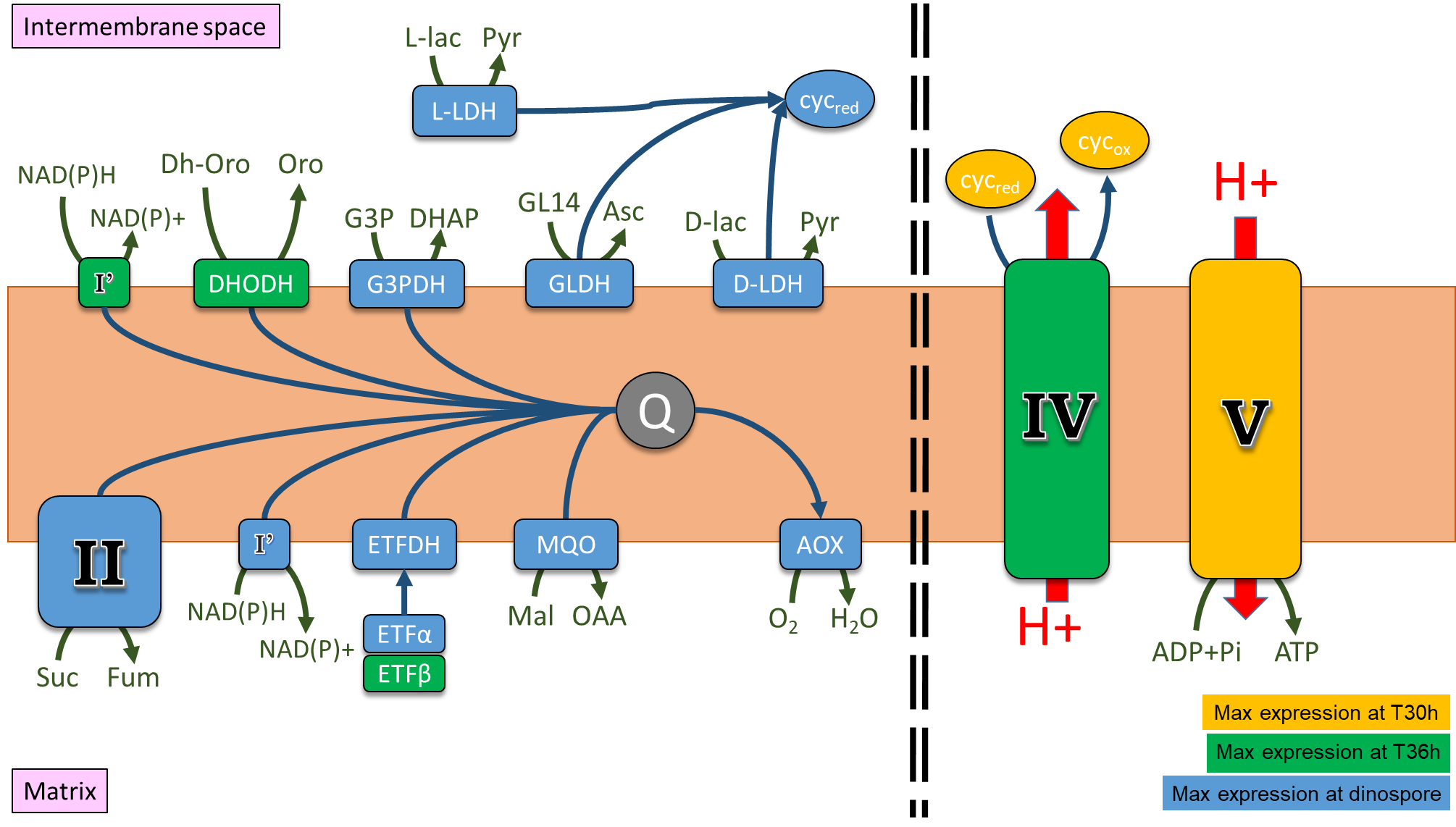

**Figure S10: Reconstruction of the Electron Transport Chain (ETC) of the Oxidative Phosphorylation (OXPHOS) pathway in the syndiniales parasite *Amoebophrya* sp. strains A120 and the expression of genes during infection.** Dashed lines illustrates the discontinuity between the ETC and proton transport in the OXPHOS pathway of *Amoebophrya*. Genes are colour-coded based on maximum expression estimated by averaging RNA-seq mapping for triplicates. Red arrows show the transfer of protons; blue arrows represent the transfer of electron. I’: NADH:ubiquinone oxidoreductase; II: succinate dehydrogenase; IV: cytochrome c oxidase; V: mitochondrial ATP synthase; AOX: alternative oxidase; cyc: cytochrome c (red and ox refer to the reduced and oxidised states); DHODH: dihydroorotate dehydrogenase; D-LDH: D-2-hydroxyglutarate dehydrogenase; ETFα and β: electron-transferring flavoproteins subunits α and β; ETFDH: electron-transferring-flavoprotein dehydrogenase; G3PDH: glycerol-3-phosphate dehydrogenase; GLDH: L-galactono-1,4-lactone dehydrogenase; L-LDH: mitochondrial cytochrome b2; MQO: malate:quinone oxidoreductase. OAA: oxaloacetate; Asc: ascorbate; D- and L-lac: D- and L-lactate; Dh-Oro: dihydroorotate; DHAP: dihydroxyacetone phosphate; Fum: fumarate; G3P: glycerol 3-phosphate; GL14: galactono-1,4-lactone; Mal: malate; Oro: orotate; Pyr: pyruvate; Suc: succinate. Q: ubiquinone pool.
